# Spatial genotypic patterns within monospecific stands of a dominant coral revealed through photogrammetry-guided sampling

**DOI:** 10.64898/2026.07.19.739434

**Authors:** Dennis van Hulten, Libby Liggins, Matan Yuval, Mary A. Sewell, Pim Bongaerts

**Affiliations:** School of Biological Sciences, The University of Auckland, Auckland, New Zealand; California Academy of Sciences, San Francisco, California, USA; Department of Collective Behaviour, Max Planck Institute of Animal Behaviour, Konstanz, Germany; Centre for the Advanced Study of Collective Behaviour, University of Konstanz, Konstanz, Germany

## Abstract

Local and global stressors are driving declines in competitive reef-building corals and favouring stress-tolerant, weedy species. In many of these taxa, rapid asexual reproductive strategies promote the proliferation of existing genotypes at small spatial scales, potentially leading to the formation of monospecific stands (“carpets”). Such reproductive strategies raise concerns for the intraspecific diversity and resilience of future reefs. The genetic structure of these spatially dominant, weedy corals remains poorly understood, in part because densely aggregated colonies lack distinct boundaries, hindering the use of standard sampling designs. We employed a photogrammetry-guided, spatially explicit sampling approach to assess the genotypic diversity and genet distribution in the increasingly dominant scleractinian coral *Madracis auretenra* across three reef sites in Curaçao. Cover of *M. auretenra* varied widely among sites and reef zones (4.1-26.2%). Genome-wide genotyping of 678 samples collected at 1-m intervals identified two genetically distinct lineages comprising 90% and 10% of the samples, with contrasting genotypic diversity (genet-to-ramet ratio Ng/N = 0.287 and 0.481, respectively). Monospecific stands (n = 20) usually contained multiple genotypes and occasionally even both lineages, but were typically dominated by a single genotype (mean 65.5%, minimum 42.3%). Simulated sampling demonstrated that combining monostands, patches, and isolated colonies would maximise captured genotypic diversity for restoration applications. Our results demonstrate that photogrammetry-guided sampling provides an effective, unbiased framework for population-genetic studies of monostand-forming corals, while revealing substantial genotypic diversity within a species that is becoming increasingly dominant on southern Caribbean reefs.

## 1 Introduction

Coral reefs are facing unprecedented declines in reef-building coral cover, posing a significant threat to the stability of these ecologically and economically important ecosystems (De’ath et al., 2012). Increasing sea surface temperature (Muniz et al., 2019), storm intensity (Gardner et al., 2005), disease outbreaks (Alvarez-Filip et al., 2022; Gladfelter, 1982), and anthropogenic disturbance (Cramer et al., 2020) are expected to continue contributing to this decline in the foreseeable future (Heron et al., 2016). However, not all coral species are equally affected by these stressors, and variations in susceptibility to local and global stressors can lead to drastic changes in coral assemblages (Chapron et al., 2022; Hughes et al., 2018). The resulting decline of previously dominant species and the rise of more tolerant or resilient taxa has been described in numerous studies (Caballero Aragón et al., 2019; Hughes et al., 2018; Loya et al., 2001; Woesik et al., 2011).

Recent models predict that, even under the most optimistic Intergovernmental Panel on Climate Change (IPCC) climate scenarios, coral communities will shift from highly diverse systems to ones dominated by fewer, less susceptible species (Kubicek et al., 2019). In accordance with these expectations, various studies have reported an increase in the relative abundance of stress-tolerant and weedy species (see Darling et al., 2012) that are more resistant to, and quick to recover from, disturbance events (Pratchett et al., 2020; Cramer et al., 2021; Darling et al., 2013; Green et al., 2008; Mudge & Bruno, 2023; Speelman et al., 2023). On some reefs, this shift has been so drastic that a coral-to-coral phase shift has been described, with weedy species completely dominating reefs (Bernardi et al., 2024). Such shifts are cause for concern, as the loss of local biodiversity and the reduction of physical and ecological traits, such as calcification rate and structural complexity, can reduce reef functionality (González-Barrios et al., 2021; Perez et al., 2025). In extreme cases, the increasing dominance of a single coral species can result in the formation of monospecific stands: contiguous patches or reef areas in which one species overwhelmingly dominates the benthic cover so that the area is visually distinguishable from the adjacent growth. Depending on the growth form of the dominant coral species, such stands are sometimes referred to as “coral carpets” (often low-relief continuous mats covering the benthic substrate, e.g. Riegl & Piller, 1999), “beds” (dense aggregations of unattached coral colonies, e.g. Chimienti et al., 2026; Keesing et al., 2026), or “thickets” (tangled clusters of branching corals creating highly complex structures, e.g. Drury et al., 2019). While monospecific stands can contribute substantially to coral cover, their ecological and genetic consequences remain poorly understood.

Weedy and stress-tolerant coral species are often characterised by low-relief, fragile colonies, high levels of inbreeding, and rapid, asexual reproduction with limited dispersal potential (Darling et al., 2012; Knowlton, 2001; Lord et al., 2023). Although some of these characteristics are favoured in restoration projects on bare or highly degraded reefs (Shaish et al., 2010), these same characteristics, combined with a shift in dominance, could lead to the formation of genetically isolated inbred coral clusters, limiting genetic variation within the population (Knowlton, 2001). For example, a recent study focused on the weedy coral *Porites asteroides* suggests that the species’ increased abundance in Florida may be driven by a few highly successful genotypes (Shilling et al., 2023). Intraspecific genetic diversity can significantly affect a species’ ability to respond to environmental stressors and provide valuable insights into the population’s potential resilience and adaptability (DeWoody et al., 2021; Pauls et al., 2013; Booy et al., 2000). These findings underscore the importance of assessing genotypic diversity at a local scale to support accurate estimates of resilience and inform effective reef management. This information is especially relevant in areas with limited natural biodiversity, where the genetic diversity captured within the few remaining species becomes increasingly critical for the stability of species-poor ecosystems and should be preserved (Crawford & Rudgers, 2013; Salo & Gustafsson, 2016).

*Madracis auretenra* (*sensu* Locke et al., 2007, following a proposed renaming of the previously synonymised *Madracis mirabillis* sensu Wells, 1973) is one of the few Caribbean coral species that remains relatively unaffected by increasing pressure from local and global stressors. Despite a recorded ∼20% average decline in Caribbean coral cover from 35% to 16% between 1970 and 2011 (Jackson et al., 2014), the relative coral cover of *M. auretenra* had increased from 8.1% to 17.6% on the reefs surrounding Curaçao and Bonaire between 1973 and 2014 (De Bakker et al., 2016). The species is known to form large monospecific stands where the dense, uninterrupted aggregation of branches completely obscures individual colony boundaries. Previous work has revealed gene flow barriers between *M. auretenra* populations within the Southern Caribbean, resulting in genetically isolated populations with high levels of inbreeding (Ballesteros-Contreras et al., 2022). Asexual reproduction through fragmentation has been documented as the primary mode of reproduction in *M. auretenra* (Bak & Engel, 1979; Bruno, 1998), although the species is classified as a semi-brooder, producing oocytes with substantial yolk reserves that may have considerable dispersal potential as planulae (Vermeij et al., 2004). Following the framework provided by Darling et al. (2012), the species exhibits “weedy” traits, including low relief, fragile colonies, rapid colonisation, and frequent asexual reproduction. The species’ predominantly asexual reproductive strategy, combined with the tendency to form monospecific stands, is expected to promote spatial clustering of genotypes and a reduction of local genetic diversity. However, despite *M. auretenra*’s increasingly dominant presence on Caribbean reefs, the genetic diversity or spatial genetic structure at a reefscape scale remains unresolved.

Studying the genetic diversity in monostand-forming corals is extremely challenging, as colony boundaries are indistinct or obscured, and monostands can cover extensive areas. While terrestrial studies have relied on *a priori* mapping (e.g., aerial and satellite imagery) to inform spatially explicit sampling (e.g., using global navigation satellite systems), such methods have been unreliable for similar studies in underwater environments, which are additionally constrained by time and safety limitations. Previous studies, therefore, employed transect lines, polar plots, or measurement tapes to select samples (Adjeroud et al., 2014b; Baums et al., 2005, 2014; Combosch & Vollmer, 2011; Drury et al., 2019). Though effective, these methods require time-intensive underwater site preparation and are limited in their scalability (Baums et al., 2005; Gorospe & Karl, 2013). With the advent of Structure-from-Motion (SfM) photogrammetry, high-resolution mapping of underwater reefscapes has become possible, and, when combined with genomic sampling, enables a more holistic understanding of the spatial genetic structure of marine organisms (Bongaerts et al., 2021). Such a “reefscape genomics” approach enables the implantation of rigorous sampling strategies within reef environments, potentially overcoming pervasive sampling biases associated with the semi-haphazard sampling (Gorospe et al., 2015; Riginos, 2015) typically applied in coral population genomic studies.

Here, we present a novel approach leveraging photogrammetric mapping to inform underwater sampling prior to collection, enabling unbiased, spatially explicit sampling at a reefscape scale (∼500-2,500 m_2_). Given the increasing importance of *M. auretenra* on Southern Caribbean reefs and the species’ tendency to form large monospecific stands, we used this species as a case study to examine the diversity and distribution of genotypes by documenting reef-wide spatial genetic structure across three locations in Curaçao. Our study provides critical insights into the molecular ecology of this dominant coral, with direct implications for reef management and conservation practices.

## 2 Methods

### 2.1 Workflow

To overcome the challenge posed by the dense aggregation of *Madracis auretenra* branches in vast, monospecific stands, we employed an *a priori,* grid-based sampling workflow that utilises three-dimensional maps generated through Structure from Motion (SfM) photogrammetry to guide sample selection. The workflow (Figure S1) follows a four-step approach: 1) Mapping, 2) Processing, 3) Collection, and 4) Geo-referencing (Figure 1). During the mapping step, thousands of overlapping images were collected and aligned for three-dimensional reconstruction using Agisoft Metashape Pro (version 2.1.2 Agisoft LLC). Next, orthomosaics, two-dimensional representations of three-dimensional models, were generated with a top-down orientation (parallel to the surface), and a virtual grid with a 1-meter spacing was projected onto the orthomosaic using QGIS (version 3.14). Branches of *M. auretenra* were annotated for collection if they were present within a radius of 0.33 m surrounding a grid point. Printouts of the annotated orthomosaics were laminated and used as maps during collection dives. Finally, using reduced-representation genomic sequencing, we determined the genotypes of the collected samples. We virtually matched these to the corresponding branches in the three-dimensional models, creating spatially referenced genotype maps.

**Figure 1:**
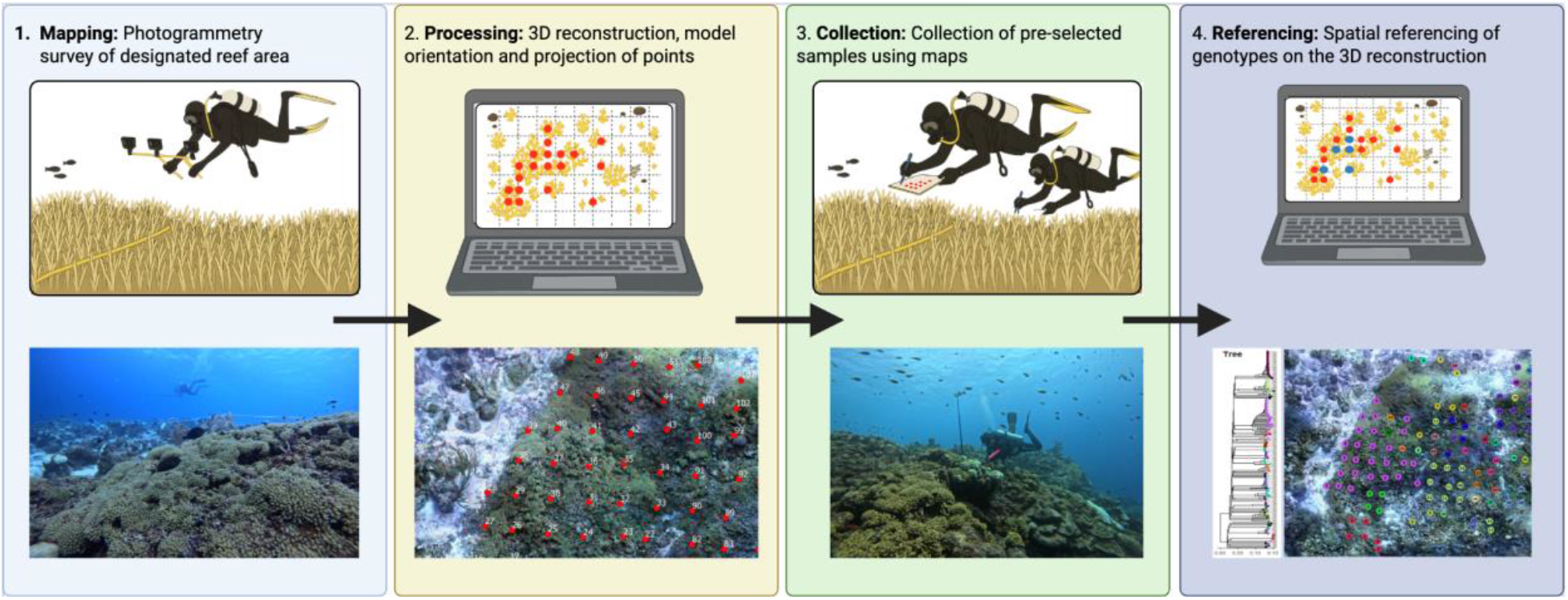
Diagram representing the photogrammetry-guided, spatially explicit sampling workflow consisting of the mapping step (1), the processing step (2), the collection step (3), and the referencing step (4).

### 2.2 Photogrammetry and site description

We collected photogrammetry data from three sites in Curaçao (Southern Caribbean): Playa Kalki, Snake Bay, and Seaquarium (Figure S2, Table S1), all located along the leeward coast and part of the “CoralScape 20K” long-term monitoring project (see Reefscape Genomics Lab, coralscape20k, 2026). The shallowest sections of Snake Bay and Playa Kalki (<4 m) comprise a sandy reef terrace environment with little coral cover. In contrast, Seaquarium is situated off an artificial peninsula where a sea wall forms an abrupt artificial boundary to the reef at a depth of 7 meters. Surveys were conducted during two field seasons: Nov 2021 - Jan 2022 and Nov - Dec 2023. During the 2021 surveys, imagery was collected with a Nikon DSLR camera equipped with four strobes, following the protocol described by Yuval et al. (2021). These initial surveys covered a transect approximately 30 m wide and continuously covered the reef between 7 and 30 m depth. Following these initial surveys, we expanded the sampling design in 2023 to include the shallow reef terrace (4 to 7 m depth) to capture most of the described depth range for *M. auretenra* (Frade et al., 2008; Vermeij et al., 2004). To increase spatial coverage per dive, we utilised a rig of three GoPro Hero 11 cameras mounted in parallel, spaced 1.5 m apart, for the 2023 surveys. Due to the site limitations at Seaquarium, no additional survey was needed to extend the area. Imagery data from each survey were processed using Agisoft Metashape Pro (version 2.1.2). The three-dimensional models proved adequate for discerning individual *M. auretenra* branches, and the resulting orthomosaics were suitable for usage as maps with a 1.5-2 mm per pixel resolution. The photogrammetry plots encompassed the 5, 10, and 20 m focal plots (100 m_2_) of the “CoralScape 20K” project at each site, so the permanently attached cattle tags could serve as depth references to orient the model’s z-axis. In addition, 50 cm aluminium bars equipped with machine-readable targets were placed on the reef to provide scaling for the plots. Both the scaling and orientation of the plots were performed in Viscore (https://chei.ucsd.edu/viscore/), using the built-in scale and orient functions. To facilitate continuity between the two surveys, we visually aligned the orthomosaics in QGIS, ensuring the grid pattern used for collections remained continuous across the two collection events.

The 2021 plots covered approximately 1,300 m² at Playa Kalki, 1,050 m² at Snake Bay, and 560 m² at Seaquarium, whereas the 2023 plots covered approximately 2,300 m² and 2,475 m² for Playa Kalki and Snake Bay, respectively. To enable area-standardised comparisons across sites, we established fixed “standardised” plots of 30 x 15 m (450 m_2_) at depths of 7-24 m. To account for variation in slope angle, its influence on structural complexity measures and the ecological relevance of reef environments on genotype distributions and coral cover, we stratified each plot into reef zones using a binned depth profile and a piecewise linear segmentation approach using *pwlf* (Jekel & Venter, 2019) to identify “breaks” in the average slope angle and divide the slope into several linear segments. Using this approach, we identified three depth-associated zones at Playa Kalki and Snake Bay, here defined as reef terrace, crest and slope. At Seaquarium, only the reef crest and slope environments were defined. Depth boundaries between zones were similar across sites, but not identical (terrace–crest transition: Playa Kalki = 5.8 m, Snake Bay = 6.4 m; crest–slope transition: Playa Kalki = 11.4 m, Snake Bay = 13.1 m, Seaquarium = 13.0 m). For cross-site comparisons, we standardised zone boundaries using mean rounded depths of 6 m (terrace–crest) and 12.5 m (crest–slope). To quantify structural complexity within reef zones, we calculated area-based rugosity as the ratio between the 2D and 3D surface area (Friedman et al., 2012) for each reef zone using the Python package *substrata* (Reefscape Genomics Lab, substrata, 2026)

### 2.3 Sample collection

We employed a minimally invasive sampling method to collect tissue samples from *M. auretenra* branches. Small coral fragments were collected using bone cutters and stored in 0.5 mL tubes filled with seawater until the end of the dive. After collection, samples were transferred to 96% ethanol at-20 ℃ prior to shipment. Some tissue samples were split during lab processing to create technical replicates, serving as a reference for clone detection during data analysis. The initial collections in 2021 covered the reef slope (7-30 m depth) at all three sites. Subsequent collections in 2023 were conducted in the shallows, continuing from the previous collections at Playa Kalki and Snake Bay; no additional samples were collected at Seaquarium. Collection events were recorded using a GoPro Hero 11 at a Point-Of-View (POV) angle to capture the sample collection process. We selected a total of 681 tissue samples for sequencing (including technical replicates), comprising 345 from Snake Bay, 109 from the Seaquarium, and 227 from Playa Kalki. We were able to position 628 collected samples within the three-dimensional plots; any remaining samples could not be mapped due to unreliable reconstruction at the edges of the surveyed area, where there is a coverage discrepancy between the orthomosaic and the 3D point cloud.

### 2.4 Molecular processing and quality control

DNA extractions and sequencing were performed by Diversity Array Technologies in Canberra, Australia. Samples were sequenced using the reduced-representation genome sequencing approach, DArTseq (Kilian et al., 2012). The resulting raw sequences are available in the NCBI Sequence Read Archive (SRA; BioProject PRJNA1475881). Quality control of the resulting raw reads was performed using FastQC (version 0.12.1) and MultiQC (version 1.27). Reads were trimmed to remove adaptors, low-quality sequences (Quality: <20, phred=33), and reads shorter than 30 bp using Trim Galore (version 0.6.10-1). The raw reads were processed using Ipryad (version 0.9.102, Eaton & Overcast, 2020) with default settings for the “ddrad” option. Reads were mapped against a reference genome for *M. auretenra* created by the Sanger Sequencing Institute (NCBI; BioProject PRJEB76086). The assembled reads were assessed for contamination using BWA (version 0.7.18-r1243-dirty) and BLASTN (version 2.16.0+). The resulting VCF file was filtered using vcftools (version 0.1.16), retaining sites genotyped in at least 80% of samples, a minor allele count (MAC) ≥ 1, and a minimum sequencing depth of 8. Samples with <10% genotyped sites were removed, retaining 678 samples and 34,466 sites for further analyses.

### 2.5 Population genomic assessment

We assessed initial population genetic structure using *STRUCTURE* (version 2.3.4, Pritchard et al., 2000) with the default recommended settings for burn-in runs (10,000) and repetitions (10), and *snapclust* (adegenet version 2.1.10). To ensure that the *STRUCTURE* and *snapclust* results were not affected by the presence of genetically identical samples, potential clones were identified and removed based on genetic similarity. Samples with genetic similarity greater than 98% were considered as belonging to the same clonal group (Figure S3). From each group, the sample with the lowest percentage of missing data was retained, while all others were disregarded. We used *Clumpp* (Jakobsson & Rosenberg, 2007) to determine the best-fitting number of populations (K) based on maximum likelihood scores for the *STRUCTURE* run. For the *snapclust* analysis, we used AIC and BIC to inform the best-fitting K for our data. The outputs of *STRUCTURE* and *snapclust* were combined with a neighbour-joining tree based on the Hamming distance to provide another visual representation of population structure, revealing a cryptic lineage within our data. As one lineage was numerically more represented than the other, we will refer to these as the major and minor lineages, respectively.

After detecting the cryptic lineage, we split the data by the *STRUCTURE* population assignments to create separate datasets for each lineage. Clonality was assessed across the full dataset and within each lineage separately to improve accuracy. We used a method adapted from Manzello et al. (2019), utilising an Identity By State (IBS) matrix generated through *ANGSD* (version htslib: 1.17) with reads filtered for a minimal mapping quality of 20, ensuring sites were genotyped for at least 70% of samples, removing duplicate and bad reads, retaining 2,164,473 out of 43,341,802 total sites. Appropriate thresholds for clonality were determined by identifying a split in the histogram of pairwise IBS distances (Figures S4) and verified using the IBS distances between replicate pairs. Because the IBS method is known to be sensitive to heterozygous SNPs (Lord et al., 2023), we also employed a clonal detection workflow based on genetic distance between individual samples to verify our results. Genetic distance was calculated based on the Hamming distance. A threshold for clone detection was set guided by the genetic distance between known replicates (98%).

To minimise biases in subsequent analyses arising from clonal samples or an imbalanced distribution of samples across the two lineages, we generated four separate data sets from our original data. The first dataset, containing all samples that passed the initial quality filtering, including clonal samples, was used to assess overall clonality and the distribution of genotypes within and among monostands. In the second dataset, clonal samples were removed, retaining only one sample per genotype selected for the highest percentage of sequenced sites. This dataset was again assessed for genetic structure using *STRUCTURE*, *snapclust*, *DAPC* (adegenet, version 2.1.11), and *PCA* (ade4, version 1.7-23) analyses. After this final assessment of population structure, we created separate datasets for each lineage, disregarding potential hybrids in these analyses. Each dataset was created from the original assembly and filtered individually using the aforementioned filtering steps. The overall genetic diversity of the population was assessed using allelic richness, F-statistics, and heterozygosity in *adegenet* (version 2.1.11), *hierfstat* (version 0.5-11), and *vegan* (version 2.7-1). These analyses were performed both within each lineage using the separate datasets and between lineages using the combined dataset to avoid overestimations of genetic differentiation due to the presence of cryptic lineages (see Sheets, 2018).

### 2.6 Reefscape genomic analysis

The physical locations of collected samples were annotated on the three-dimensional plots in Viscore using POV collection footage. To calculate cover percentages and map aggregations, we traced *M. auretenra* branches in the orthomosaics using Agisoft Metashape Pro’s built-in draw polygon tool. We measured their size using the Python package *shapely* (version 2.1.2). We defined monospecific stands as continuous aggregations occupying an area greater than *π* m_2_, following the largest reported colony size in the forereef environment of Curaçao (up to 2 m in diameter; Bruno, 1998). In addition, we identified “patches”, aggregations not big enough to be classified as monospecific stands (area between 1 and *π* m_2_), and isolated colonies (area <1 m_2_). Due to differences in stand shape, the number of samples per monospecific stand varied widely. To ensure our genetic comparison between stands was not influenced by extremely small sample sizes, we included only monospecific stands with five or more samples in the genetic analysis. Our data comprised 262 samples from monospecific stands, 187 from patches, and 156 individual colonies. The distance between samples was calculated using the substrata package. Leveraging photogrammetry maps and genomic data from our initial collections (2021), we revisited previously sampled colonies during the 2023 survey after the discovery of a cryptic lineage to gather underwater macro photographs and skeletal fragments from a subsample of each lineage (n = 7 per lineage).

## 3 Results

### 3.1 Demographic analysis of photogrammetry data

The overall cover, calculated as the total area of the traced polygon of *Madracis auretenra* colonies, varied between sites. We found higher overall cover at Snake Bay, with colonies covering 172.2 m_2_ of the approximately 2,475.0 m_2_ of reef surveyed (6.9%), and Seaquarium 42.0/560.0 m_2_ (7.5%) compared to Playa Kalki 84.8/2,300.0 m_2_ (3.7%) (Figure 2A, Table 1). *M. auretenra* cover was unequally distributed between reef zones, with the highest cover found at the reef crest (∼6.0-12.5 m) in Playa Kalki (∼65.6% of all *M. auretenra* cover) and Snake Bay (58.6% of all *M. auretenra* cover) (Table 1). In contrast, most of the *M. auretenra* cover at Seaquarium was found at the reef slope (<12.5 m; 88.1% of all *M. auretenra* cover), though the physical limitations of this site restrict comparisons with the other sites. When considering the standardised plots (450 m_2_; only containing reef crest and slope), *M. auretenra* was decidedly more dominant at Snake Bay (117.9/450.0 m_2_, 26.2%) compared to Seaquarium (38.3/450.0 m_2_, 8.5%) and Playa Kalki (18.4/450.0 m_2_, 4.1%) (Figure 2A). We identified 20 monospecific stands across the three sites. We found considerably more monospecific stands at Snake Bay (n = 12) compared to Playa Kalki (n = 5) or Seaquarium (n = 3) (Figure 2B). Inside the standardised areas, we found 8 stands at Snake Bay, 2 at Seaquarium, but none at Playa Kalki. Monospecific stands across all sites had a mean perimeter of 26.2 m (SD = 16.1 m) and a mean spatial extent of 10.4 m_2_ (SD = 9.1 m_2_). The mean stand area was roughly 1.65× larger at Snake Bay (mean ± SD = 12.1 ± 10.7 m_2_) compared to Playa Kalki (mean ± SD = 7.3 ± 5.0 m_2_) and Seaquarium (mean ± SD = 8.3 ± 8.0 m_2_); however, this difference was not statistically significant (Gamma GLM, χ^2^ = 0.98, p = 0.51). The largest monospecific stand was described at Snake Bay, reaching a spatial extent of 40.0 m_2_ and a perimeter of 53.4 m.

**Figure 2:**
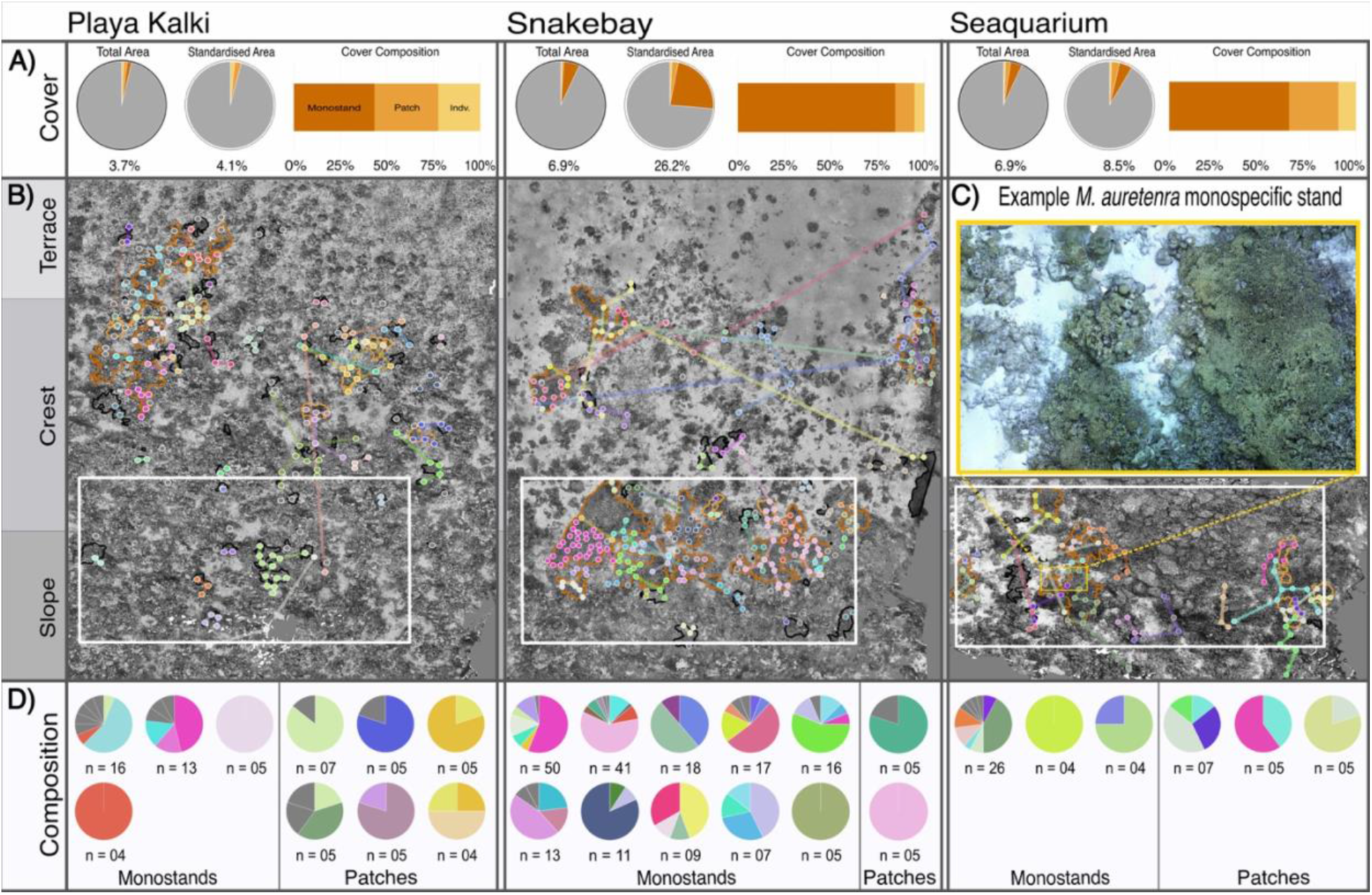
Visual overview combining photogrammetry and genomic data. A) the overall cover of *M. auretenra* colonies in the full and standardised area (450 m2 between 7 and 24 m depth). Colours represent monospecific stands (dark orange), patches (orange), and individual colonies (yellow). B) Distribution of genotypes at Playa Kalki, Snake Bay, and Seaquarium. Colours represent genets, and lines connect all ramets within a genet. Monospecific stands are highlighted in dark orange, and patches are highlighted in orange. The white box indicates the standardised area. The yellow box highlights an example of the dense aggregation of *M. auretenra* branches in a monospecific stand at Seaquarium. C) Visual representation of the genotypic composition within monospecific stands and patches. Each pie chart represents a single stand or patch, with the n value indicating the number of samples collected from that stand or patch. The colours represent the different genotypes, with dark grey indicating unique genotypes. These colours match those presented in B).

**Table 1:** Summary of demographic measurements of all monospecific stands (n=20) divided per site and reef zone as measured in the total surveyed area at each site. Stands that span multiple reef zones are included in all relevant zones.

| *Site* | *reef zone* | % *M. auretenra* cover of total area | # Monostands | Monostand extent avg (m<sup>2</sup>) | Monostand extent max (m<sup>2</sup>) | Monostand perimeter avg (m) | Monostand perimeter max (m) | *Structural rugosity* |
| --- | --- | --- | --- | --- | --- | --- | --- | --- |
| *Playa Kalki* | *Slope* | 0.3 | 0 | - | - | - | - | *1.785* |
| *Playa Kalki* | *Crest* | 1.8 | 5 | 7.34 | 13.04 | 27.51 | 44.99 | *1.784* |
| *Playa Kalki* | *Terrace* | 0.6 | 3 | 6.99 | 13.04 | 26.63 | 44.99 | *1.401* |
| *Snake Bay* | *Slope* | 11.6 | 7 | 14.44 | 39.98 | 29.05 | 57.10 | *1.703* |
| *Snake Bay* | *Crest* | 11.2 | 11 | 12.90 | 39.98 | 26.59 | 57.10 | *1.479* |
| *Snake Bay* | *Terrace* | 0.2 | 2 | 12.63 | 15.54 | 30.11 | 33.16 | *1.282* |
| *Seaquarium* | *Slope* | 7.8 | 3 | 8.31 | 17.51 | 26.67 | 55.56652 | *1.791* |
| *Seaquarium* | *Crest* | 2.5 | 2 | 10.81 | 17.51 | 34.57 | 55.56652 | *1.718* |
| *Playa Kalki* | *standardised* | 4.1 | 0 | - | - | - | - | *1.932* |
| *Snake Bay* | *standardised* | 26.2 | 7 | 14.4 | 39.9 | 28.9 | 57.0 | *1.677* |
| *Seaquarium* | *standardised* | 8.5 | 2 | 10.8 | 17.5 | 33.7 | 54.0 | *1.785* |

Monospecific stands of *M. auretenra* at Snake Bay covered the largest combined area on the reef in both the full plots (Snake Bay = 145.4 m_2_, Playa Kalki = 36.7 m_2_, Seaquarium = 24.9 m_2_) and in the standardised plots (Snake Bay = 105.0 m_2_, Playa Kalki = 0 m_2_, Seaquarium = 21.6 m_2_) (Figure 2A, Table 1). The abundance and spatial extent of individual colonies (<1 m_2_) and patches (1 to *π* m_2_) varied between sites. Within the standardised plots, more individual colonies were found at Playa Kalki (n = 49, total cover = 7.5 m_2_) and Seaquarium (n = 51, total cover = 3.3 m_2_) compared to Snake Bay (n = 25, total cover = 3.3 m_2_). The abundance of patches within the standardised area was comparable across all three sites: 6 patches covering 10.9 m_2_ at Playa Kalki, 5 covering 10.6 m_2_ at Snake Bay, and 5 covering 13.4 m_2_ at Seaquarium. Outside of the standardised plots, we find decidedly more individual colonies and patches at Playa Kalki (n = 92 and n = 17) covering a decidedly larger area (19.1 m_2_ and 29.0 m_2_) compared to Snake Bay (n = 65 and n = 9 covering 9.36 m_2_ and 17.8 m_2_).

### 3.2 Prevalence of asexual reproduction and distribution of genotypes

Our analysis of the sequencing data revealed two genetically dissimilar clusters with dissimilar sample sizes (n=613, n=65) (See section 3.3). After detecting these two lineages, we split the data by cluster for further analysis. Following the thresholds defined by comparing IBS distances as node heights (0.0156 and 0.018; Figure 3A, Figure S3A, B), less than a third of the samples exhibited unique genotypes (204/681, Ng/N = 0.299). Our analysis of the IBS matrices for the full data set and each genetic cluster separately revealed clear clonal groupings that were consistent across the full data analysis and the separate data sets (Figure 3B, Figure S3C, D). Genotypic diversity, measured as the genet-to-ramet ratio (Ng/N), differed between lineages, with the major lineage exhibiting lower diversity (176 unique genotypes among 614 samples; Ng/N = 0.287) than the minor lineage (28 unique genotypes among 65 samples; Ng/N = 0.431) (Table 2). A binomial generalised linear model revealed a significant positive association between genotypic diversity and structural rugosity after accounting for reef zone and lineage (β = 2.27 ± 0.76 SE, p = 0.003; Table S1). Reef zone also significantly influenced genotypic diversity independent of rugosity, with lower Ng/N ratios on the reef slope (β = - 0.57 ± 0.21, p = 0.007) and higher Ng/N ratios on the reef terrace (β = 0.77 ± 0.36, p = 0.031) relative to the reef crest. No evidence of overdispersion was detected (dispersion ratio = 1.01). Lineage remained a significant predictor of genotypic diversity after accounting for rugosity and reef zone (β = 0.57 ± 0.29, p = 0.047).

**Figure 3:**
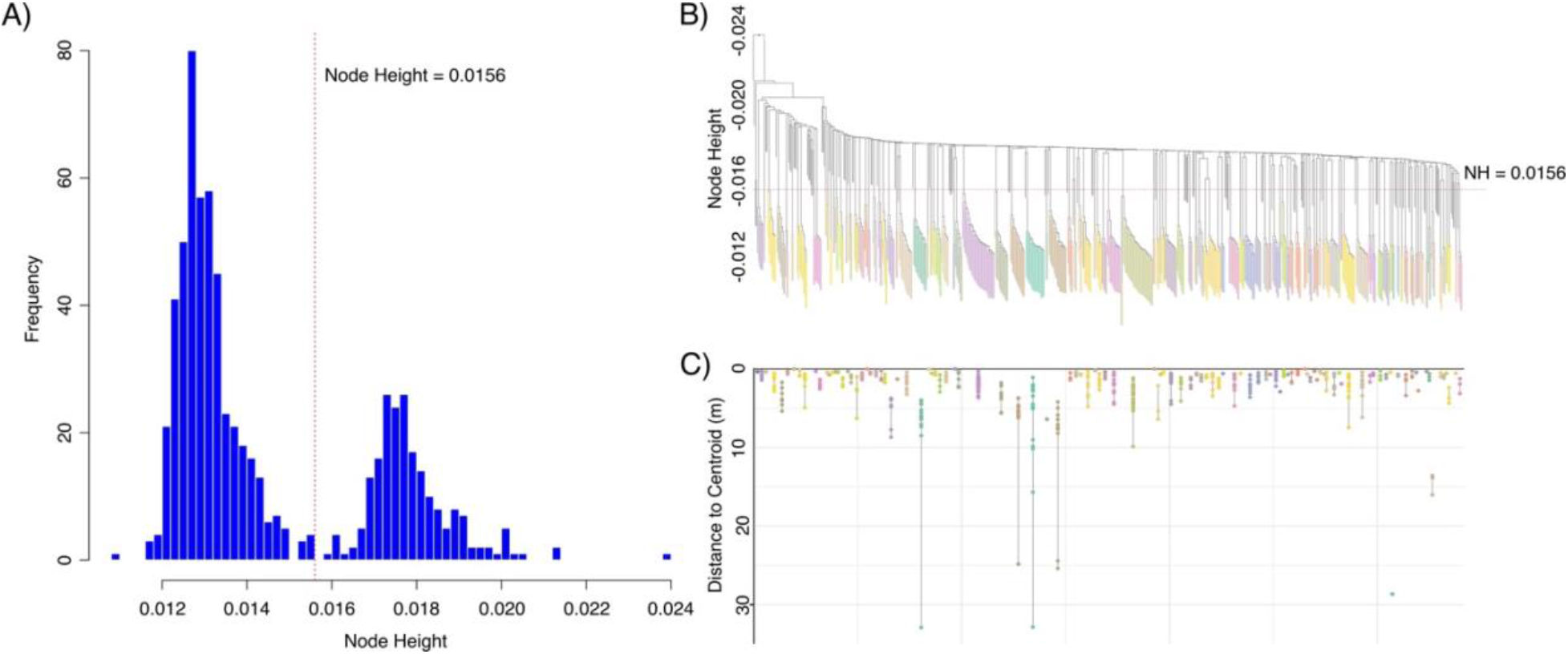
Results of the Identity by State distance analysis used to inform clonal groups. A) Histogram showing frequencies of node heights based on IBS distance, the red dotted line indicates the threshold selected to inform clonal groups. B) Dendrogram based on IBS distance with clonal groups indicated by colour and unique genotypes highlighted in dark grey. C) A visual representation of physical clustering of clones with the distance of each individual from the clonal group’s “centroid” calculated as the average x, y, z coordinates of the group.

**Table 2:** A summary of genotypic diversity measured as the genet-to-ramet ratios across lineages, sites and reef zones.

| Genet-Ramet ratio (Ng/N) | Snake Bay | Playa Kalki | Seaquarium | Across sites |
| --- | --- | --- | --- | --- |
| major lineage | 70/300 = 0.232 | 85/227 = 0.374 | 26/87 = 0.299 | 176/614 = 0.287 |
| Reef terrace major lineage | 10/36 = 0.278 | 15/32 = 0.469 | NA | 25/68 = 0.368 |
| Reef crest major lineage | 44/149 = 0.295 | 61/143 = 0.427 | 2/3 = 0.667 | 107/295 = 0.363 |
| Reef slope major lineage | 21/82 = 0.256 | 10/30 = 0.333 | 21/79 = 0.266 | 53/192 = 0.276 |
| minor lineage | 17/39 = 0.436 | 4/5 = 0.800 | 6/21 = 0.286 | 28/65 = 0.431 |
| Reef terrace<br>minor lineage | NA | 1/1 = 1 | NA | 1/1 = 1 |
| Reef crest<br>minor lineage | 2/12 = 0.167 | 1/1 = 1 | 2/5 = 0.4 | 5/18 = 0.278 |
| Reef slope<br>minor lineage | 12/23 = 0.522 | 2/3 = 0.667 | 5/14 = 0.357 | 20/40 = 0.500 |

Clonal group size, measured as the number of ramets per genet, varied greatly between sites. The two largest clonal groups were found at Snake Bay, each comprising 31 samples. In total, we identified 12 clonal groups (genets) comprising over 10 individual samples (ramets). The majority of these large clonal groups were found in Snake Bay (8/12). Clonal group size differed significantly among sites, with Snake Bay supporting slightly larger clonal groups (mean number of ramets ± SD = 6.09 ± 6.51) compared to Playa Kalki (mean ± SD = 4.05 ± 2.97) and Seaquarium (mean ± SD = 4.65 ± 2.50) (negative binomial GLM estimate = 0.39 ± 0.17 SE, p = 0.021). However, this contrast was marginal after Tukey adjustment through a pairwise Estimate Marginal Means test (p = 0.05, 0.06). Clonal group sizes did not differ between Playa Kalki and Seaquarium (negative binomial GLM estimate =-0.04 ± 0.21 SE, p = 0.823). The average physical distance between clone pairs across all sites was 3.16 ± 0.62 meters, with no significant differences between sites (Kruskal-Wallis χ^2^ = 1.02, df = 2, p-value = 0.601). Although most ramets of the same genet seemed to cluster together at relatively short distances, the maximum distance between clonal samples far exceeded the average distance, reaching up to 39.94 meters at Snake Bay, with maximum distances of 23.59 m and 11.14 m at Playa Kalki and the Seaquarium, respectively (Figure 2C, 3C).

To ensure adequate sample sizes for genetic analyses, we selected monospecific stands (area > *π* m_2_) with at least 5 samples for comparison of genetic and genotypic diversity, resulting in the evaluation of 18 monospecific stands. From those 18 stands, 14 contained more than one genotype, with up to 9 genotypes recorded in a single monospecific stand (Figure 2C). In the vast majority (n = 13) of stands, a single genotype accounted for over 50% of the sequenced samples (mean = 65.5%). In the remaining five stands, the most common genotype still accounted for at least 42.3% of the sequenced samples. A similar pattern was observed in patches (area < *π* m_2_), with 10 out of 11 patches containing more than 1 genotype. Similar to monospecific stands, patches were typically dominated by a single genotype, with 9 of 11 patches containing at least 50% ramets of the same genet (Figure 2C).

Genotypic diversity was lower in monospecific stands across all sites (Ng/N = 0.267) compared to isolated colonies (Ng/N = 0.564) and patches (Ng/N = 0.417). The genotypic diversity per stand, quantified as genet-to-ramet ratios (Ng/N), varied among stands, ranging from 0.167 to 0.800 (mean ± SD = 0.374 ± 0.151), but did not differ significantly among sites (Kruskal-Wallis test, χ^2^ = 0.77, df = 2, p = 0.68). Stand size had a significant positive effect on the number of genotypes found within the stand (Pearson’s product-moment correlation, df = 236, p < 2.2 × 10⁻¹⁶, cor = 0.83). Genotypes were occasionally shared between monospecific stands, with 4.4% of genotypes found in multiple stands (Figure 2B). Genotypes were also shared between monospecific stands and isolated colonies (8.7%) and between monospecific stands and patches (7.6%).

We simulated genetic sampling of 20 colonies by randomly drawing individual samples from either monospecific stands, patches, or isolated colonies over 100,000 iterations. This simulation revealed that sampling within individual colonies yielded a higher average number of unique genotypes (18.2) than within patches (17.5) or monospecific stands (15.5). In addition, sampling from individual colonies had a higher probability of capturing all unique genotypes than in monospecific stands and patches (isolated colonies: 12.6%; patches: 5.6%; monospecific stands: 0.3%). When considering sampling from a combination of isolated colonies, patches, and monospecific stands, we found that including isolated colonies tended to increase the number of unique genotypes and including monospecific stands tended to decrease it (Table S4). Repeating this simulation but only considering sampling within a single site revealed that, with the exception of individual colonies at Playa Kalki, sampling within a combination of monospecific stands, isolated colonies, and patches yielded the highest average number of genotypes (Table S5).

### 3.3 Cryptic genetic lineages with asymmetrical depth distribution

Our data revealed two genetically distinct lineages of *M. auretenra* in Curaçao, which were consistently supported by a neighbour-joining analysis (Figure 4A), principal component analysis (PCA) (Figure S4), a DAPC (Figure 4B), and STRUCTURE results (Figure 4C, S5). One lineage was numerically more abundant (n = 613, the “major lineage”) compared to the other (n = 65, the “minor lineage”). Our initial analyses revealed moderate genetic differentiation between the lineages (F_ST_ = 0.17, Amova ɸ_st_ = 0.256, simulated p-value = 0.001), with approximately 3% of SNPs (1,206/34,466) exhibiting private alleles exclusive to a single lineage. We found no alternatively fixed alleles. Although we observed little evidence of admixture between the lineages, one potential hybrid between the major and minor lineages showed roughly equal assignment to each lineage in the STRUCTURE results. This assignment coincided with the location of the sample in both PCA and DAPC analyses and aligned with the position of the sample in the neighbour-joining tree (Figure 4A, B and S4). The DAPC analysis identified 4 additional potential hybrid samples (Figure 4B). However, they were not detected in the STRUCTURE or neighbour-joining trees (Figure 4A, C) and were therefore not classified as hybrids in follow-up analyses.

**Figure 4:**
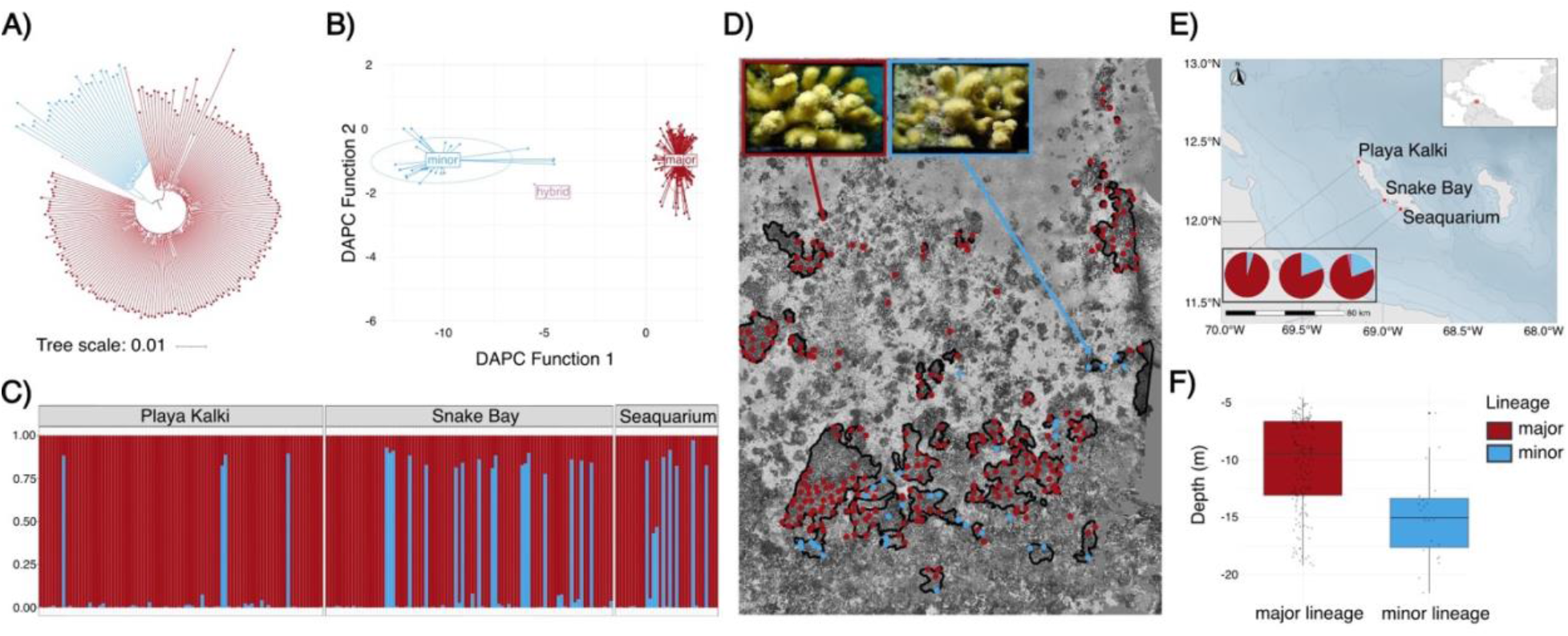
Overview of the divergent genetic lineages. A) Midpoint rooted Neighbour joining tree based on Hamming distance of the minor (blue) and major (red) lineages, in combination with the presence of a putative hybrid (purple). B) DAPC results showing the two lineages and 4 putative hybrids. C) Structure results for K=2 with samples grouped by site. D) Overview of lineage distribution at Snake Bay, highlighting the co-occurrence of lineage within monospecific stands, macrophotographs of two individual representatives from the minor (blue) and major (red) lineages. E) The relative abundance of the major and minor lineages across the sampling sites. F) A boxplot showing the depth distribution across all sites of individual samples separated by lineage.

Using the dataset without clonal samples and separated by genetic lineage (minor lineage n = 27, major genotype n = 176), we found no genetic differentiation between sites (Fst < 0.0001 for both lineages) which was supported by an Isolation by Distance (IBD) analysis, comparing genetic distance measured in Hamming distance with geographical spacing between sites (Mantel r =-0.005, p-value = 0.61). Calculated separately for each lineage by splitting the commonly filtered dataset, the inbreeding coefficient was higher in the minor lineage than in the major lineage (F_is_ = 0.250 vs 0.142, respectively). In addition, we found that expected and observed heterozygosity were higher in the minor lineage (Hs = 0.216, Ho = 0.171) than in the major lineage (Hs = 0.152, Ho = 0.130).

After the detection of the two cryptic lineages, macrophotos and skeletal fragments from each lineage were collected for representative samples (n = 14). Samples from both lineages conformed to the original description of *Madracis auretenra* by (Locke et al., 2007). An initial informal comparison, focussing on corallite density, spacing and number of septa revealed no obvious differentiating traits (Figure S12). In terms of spatial distribution, representatives from different lineages often occurred in close physical proximity on the reef, with 7 monospecific stands and 4 patches containing samples from both lineages (Figure 4D). Representatives of both lineages were present at all three sites; however, the minor lineage was less abundant at Playa Kalki than at the other two sites (Figure 4E). Depth distributions differed significantly between lineages when pooled across all sites (Wilcoxon rank-sum test, p-value = 6.82 × 10_-6_), with major lineage samples spanning the full sampled depth range (∼4-22 m, mean ± SD = 10.1 ± 3.9 m). In contrast, the minor lineage appeared virtually absent from the shallow reef terrace (mean ± SD = 14.6 ± 3.8 m) (Figure 4F, S9, S10, S11). Lineage-specific depth distributions differed significantly between sites (Kruskal-Wallis test, p-value = 1.372 × 10_-13_; Figure S8). A linear model including site and lineage revealed a significant site-by-lineage interaction (Lineage x Site: *F (2, 174)* = 7.46, p = 0.0008) with post-hoc comparisons showing significant depth differentiation at Snake Bay (emmeans p-value < 0.0001) but not at Playa Kalki (p = 0.28) or Seaquarium (p = 0.49).

## 4 Discussion

In this study, we demonstrate how high-resolution SfM photogrammetry can be leveraged to allow for unbiased, spatially explicit genomic sampling of monostand-forming marine invertebrates. Applying this method to the weedy coral *Madracis auretenra,* we document that despite the species’ predominant asexual reproductive strategy, the populations harbour moderate genotypic diversity. Our data show that a single genotype typically dominates large aggregations (monospecific stands or patches); however, the majority harbour additional, lower-abundance genotypes. We discuss how these patterns can guide ramet selection to maximise genotypic diversity in coral restoration interventions. We found no evidence of genetic separation over the studied geographical distances. Still, we document two morphologically similar yet genetically divergent lineages with overlapping depth distributions and limited signs of admixture, suggesting some degree of reproductive isolation. We discuss the effectiveness of our methodology for documenting genotypic diversity and distribution, and explore the implications of our findings for the general understanding of the patterns driving genotypic diversity in *M. auretenra* populations.

### 4.1 Demographic analysis of photogrammetry data

Haphazard or spatially clustered sampling has long been the predominant approach in marine population genetic studies, although this can potentially introduce biases, particularly in systems dominated by asexual reproduction (Gorospe et al., 2015; Riginos, 2015). While truly random or spatially comprehensive sampling regimes provide the most reliable data (Gorospe et al., 2015), such approaches are resource-intensive and hindered by the logistical constraints of working underwater. Previous studies have utilised transect-based approaches to achieve spatially comprehensive sampling within dense aggregations (e.g. Drury et al., 2019) or polar plots to achieve random sampling (Baums et al., 2006). Though effective, these methods require extensive setup time underwater and are therefore difficult to conduct over wide depth ranges or large areas. In this study, we leverage SfM photogrammetry to approach spatially comprehensive sampling in a system where individual colony boundaries are obscured, and a single 1 m_2_ patch can contain >4,000 branches (Bruno, 1998). By implementing a grid-based sampling regime encompassing both aggregations (n = 50) and individual colonies (n = 261) across three sites, we endeavoured to minimise bias while achieving a sample size large enough to approximate accurate genotypic diversity (n = 678). The use of photogrammetric maps allowed us to shift the preparation for spatially explicit sampling from underwater to a desktop exercise, making the underwater sampling effort more similar to that of haphazard sampling regimes. This approach may also be relevant to other marine ecosystems that exhibit strong dominance by particular species (e.g., mussel reefs, seagrass meadows, and sponge gardens) and where comprehensive sampling is unfeasible, as individuals cannot be clearly delineated.

Characterised by a mainly heterotrophic feeding strategy (Sebens et al., 1996), quick growth through fragmentation (Brito-Millán et al., 2019) and highly diverse microbial communities (Wallace et al., 2025), the weedy coral *M. aurentenra* has become increasingly dominant on Curaçao (De Bakker et al., 2016). Our data further demonstrate this dominance at a reefscape scale. Branches of *M. auretenra* covered up to 6.9% of the surveyed area in Snake Bay, with most of the cover accounted for by large monospecific stands reaching areas of up to 40 m_2_. Monospecific stands accounted for the largest portion of total *M. auretenra* cover across all sampled sites, highlighting the importance of monospecific growth for *M. auretenra* colonies and potentially indicating self-facilitation or positive density dependence. For instance, in shallow reef environments characterised by high irradiance and flow conditions, high branch densities may promote nutrient uptake by reducing flow speed and alleviating irradiance stress from self-shading (Sebens et al., 1997). In addition, the branching morphology of this species appears to adapt to flow conditions, creating a stagnant region in high-flow conditions that could promote prey capture (Kaandorp et al., 2003). The ability to alter the flow conditions around the colony has been hypothesised to contribute to the increased diversity of microbial communities observed in this species (Wallace et al., 2025). Collectively, these findings suggest that the dense aggregation of branches in monospecific stands may represent a key ecological strategy underlying the increasing dominance of *M. auretenra* on Curaçao’s reefs.

The total cover of *M. auretenra* varied greatly across the three sites when considering the reef crest and slope (in the standardised areas) from 26% at Snake Bay to 4% at Playa Kalki, and appears negatively correlated with structural complexity. Previous work on Curaçao suggested higher growth and per cent coverage of *M. auretenra* in sites with higher anthropogenic nutrient loading (Cook, 2022). Although we did not measure nutrient loading, our results are comparable to those presented by Cook (2022) across the same or similar sites, with higher *M. auretenra* cover recorded at Seaquarium, while the less nutrient-loaded Playa Kalki shows the lowest overall cover. However, despite being located in an area with less nutrient loading, Snake Bay showed the highest overall cover of *M. auretenra,* indicating that nutrient loading alone does not fully explain the prevalence of *M. auretenra*, and other site-specific characteristics might drive the dominance of the species. Variation of coral cover and colony density within monospecific coral stands across reef zones has previously been linked to hydrodynamic regimes (Vargas-Ángel et al., 2003). Coral species highly adapted to fragmentation, such as *M. auretenra* (Bruno, 1998; Nagelkerken et al., 2000), are predicted to become increasingly dominant in high-wave-energy environments, where their life-history traits facilitate clonal proliferation while less resilient competitors suffer significant colony damage (Highsmith, 1982). A recent example of this process has been reported for the hydrozoan *Millerpora* on the reefs of Mo’orea (Dubé et al., 2017). In addition, hydrodynamic force models and observational data have shown that wave energy exposure can drive the dominance of certain coral species over others (Storlazzi et al., 2005). Snake Bay is exposed to the highest wave energy, followed by Seaquarium and lastly Playa Kalki with minimal exposure (Van Duyl, 1985). Perhaps this could explain variation across reef zones, with the highest percentage cover of *M. auretenra* at both Playa Kalki and Snake Bay within the reef crest environment, where wave energy is the highest (Van Duyl, 1985; Ferrario et al., 2014; Kench et al., 2022).

4.2 The prevalence of asexual reproduction on monospecific stands of *Madracis auretenra*

Despite the potential for extensive asexual reproduction through fragmentation described for *M. auretenra,* our analysis of 678 samples collected from 3 sites revealed moderate genotypic diversity with an overall genet-to-ramet ratio of 0.299 (Table 2). The prevalence of asexual reproduction, as inferred from genet-to-ramet ratios, varied among sites, potentially indicating site-specific differences in fragmentation intensity or fragment survival, as previously reported for *M. auretenra* populations (Bruno, 1998). Our data revealed a positive association between structural rugosity and genotypic diversity, indicating that more rugose environments were more likely to host unique genotypes. Higher structural complexity could increase the availability of suitable habitat, promoting the settlement and survival of coral recruits and thereby enhancing genotypic diversity, similar to results found for *Pocilloporidae*, *Acropora*, *Millepora*, and other families (Fujiwara et al., 2022). In addition, greater habitat heterogeneity might result in a broader niche space, potentially supporting a more diverse range of genotypes. Alternatively, higher structural complexity could limit fragment dispersal, hampering the spread of genotypes across large areas.

On the other hand, site-specific characteristics not measured in this study but potentially correlated with structural complexity also could influence genotypic diversity. For example, exposure to higher wave energy could increase fragmentation frequency, promoting asexual reproduction in *M. auretenra* (Bruno, 1998), while simultaneously affecting the structural complexity of the environment by reducing large coral structures (Xu et al., 2025). Similarly, the prevalence of sandy substrate would greatly influence the environment’s structural complexity while hindering sexual recruitment. Although fragment survival rates in sandy substrates are severely limited (Heyward & Collins, 1985), unattached fragments of *M. auretenra* show above-average survival rates compared to other coral species (Bruno, 1998; Mercado-Molina et al., 2014; Nagelkerken et al., 2000), with some survival in sediment-rich environments. *M. auretenra* fragments often consist of dense branch clusters originating from a single base. Such a cluster could favour the survival of the uppermost branches that remain raised above the sandy substrate, while the lower branches provide a hard substrate to anchor the fragment in the moving sediment.

Although our results indicate moderate population-wide genotypic diversity, aggregations of *M. auretenra* are typically dominated by a single genotype. Such single-genotype dominance may arise from individual expansion following fragmentation, eventually leading to the gradual incorporation of adjacent colonies and the creation of a larger monospecific stand (Brito-Millán et al., 2019). Alternatively, aggregations might form through multiple sexually recruited genotypes that, through competition and/or selection, shift toward the dominance of a single genotype (Stuefer et al., 2009; Koppitz & Kuhl, 2000). Finally, the presence of a well-established adult colony might promote conspecific larval settlement, enhancing genotypic diversity around established monospecific stands (Sims et al., 2023; Vermeij, 2005). Although the processes promoting disproportional genotype dominance within monospecific stands remain unresolved, the resulting genotypic structure may enhance the ecological resilience of these stands, in line with the “insurance hypothesis” (Loreau et al., 2021; Tilman et al., 1997). Specific genotypes may be disproportionately susceptible to bleaching (Dilworth et al., 2024; Drury & Lirman, 2021; Edmunds, 1994), predation (Rivas et al., 2021) or disease outbreaks (Brown et al., 2022; K. R. Eaton et al., 2025). As such, populations comprising a single genotype may be disproportionately affected by disturbances and generally lack the resilience often associated with genotypic diversity (Ayre & Hughes, 2004). Therefore, the genotypic heterogeneity of these stands may retain ecological resilience through diversity, while maintaining the ecological integrity and competitive advantage of the stand through the dominant genotype under prevailing environmental conditions (Aspinwall et al., 2011; Hoeber et al., 2018; Münzbergová et al., 2009).

Accounting for the prevalence of asexual propagation within natural populations is increasingly critical to resilience forecasting, diversity estimations, and conservation efforts (Baums, 2008; Meloni et al., 2013; Roberts et al., 2017; Tierney et al., 2020). The inclusion of ramets in population genetic studies can affect statistical power, violate assumptions of certain analyses, or invalidate their interpretation unless precautionary steps are taken (Grünwald et al., 2017; but see Meirmans, 2025). By collecting samples in a spatially explicit manner, we endeavoured to include both clonal and unique genotypes to better understand the total genotypic diversity of *M. auretenra* populations within a reef at a specific spatial scale. As a result, our spatially explicit sampling design provides a valuable framework for informing future sampling strategies to maximise the recovery of unique genotypes while minimising redundant sampling of clonemates. Through a simulation exercise, we show that sampling individual colonies (<1 m_2_) yielded a ∼15% increase in recovered unique genotypes compared to monospecific stands and a ∼4% increase compared to sampling patches. Sampling individual colonies or a mixed approach of sampling stands, patches, and individual colonies yielded comparable numbers of unique genotypes. Combining these results with our findings on the average distance between ramets at 3.2 m, we advocate for sampling designs that ensure a minimum distance of ∼4 m between samples, focusing on collecting samples from a mixture of individual colonies, patches, and monospecific stands but avoiding sampling multiple times within a single monospecific stand.

### 4.3 Genetically divergent but cooccuring lineages

We detected two morphologically similar but genetically distinct lineages in the *M. auretenra* populations on Curaçao. Although we did not observe any obvious differentiating morphological traits between the two lineages in branch thickness, corallite density, or tissue colour, we note that the species is known to exhibit environmentally driven phenotypic plasticity between clone mates (Bruno & Edmunds, 1997). In reef environments where the lineages overlap, representatives of both lineages frequently co-occurred in a single monostand, demonstrating sympatry and the absence of physical barriers to gene flow. Despite their physical proximity and only moderate genetic differentiation, there were few indications of potential admixture between the lineages (only a single putative F1 hybrid), supportive of a certain extent of reproductive isolation. The existence of two lineages occurring in sympatry, showing genome-wide divergence and an almost complete lack of admixture, is consistent with their recognition as separate species under the genotypic cluster concept of Mallet (1995, 2020). The documentation of these cryptic taxa is timely and relevant to both scientific and conservation efforts, as cryptic diversity can bias ecological studies and affect estimates of biological and genetic diversity (Cheng et al., 2025; Chenuil et al., 2019; Grupstra et al., 2024; Riginos et al., 2024). Further studies should explore the presence of morphological, ecological or physiological differences between the two lineages reported in this study, and evaluate the potential contribution of differences in reproductive timing to the observed reproductive isolation as observed in other cryptic coral taxa (Bongaerts et al., 2021; Chamberland et al., 2025; Ricardo et al., 2025; Rosser, 2015).

The minor lineage presented in this study could represent a relatively recent migration to Curaçao. This hypothesis would be supported by the lower overall abundance of the minor lineage across our data, combined with the higher expected and observed heterozygosity in that lineage, which could suggest its origin in a larger, older population. Although we have no way to infer the potential origin of the lineage with regard to the wider Caribbean region, a previous study of *M. auretenra* populations within the Southern Caribbean region alluded to potential migration between Colombia (Punta Bota) and Curaçao, and gene flow among Barbados, Curaçao and Colombia (Bolivar-Cordoba) following population assignments from *GENODIVE* (Meirmans & Van Tienderen, 2004) and migration patterns from *divMigrate* (Sundqvist et al., 2016). However, these findings were presented as exploratory given the uncertainty of the migration estimates produced by these tools (Ballesteros-Contreras et al., 2022). Recent migration into Curaçao could explain the higher genet-to-ramet ratios observed in the minor lineage, as these genotypes would have had less time to establish compared to those in the major lineage. In addition, we note that the general current directions around Curaçao flow northwest (Bertoncelj et al., 2025), potentially explaining the lower prevalence of the minor lineage at Playa Kalki. However, as we did not sample other Caribbean locations in this study, we are unable to evaluate a potential migration origin, leaving the geographic context of diversification within *M. auretenra* for future studies.

## 5 Conclusion

Understanding the molecular ecology of weedy, monospecific stand-forming corals is increasingly relevant as environmental disturbances continue to shift reef communities toward those dominated by resilient species capable of rapidly colonising newly available space. Here, we demonstrated how Structure-from-Motion (SfM) photogrammetry can facilitate the implementation of rigorous, spatially explicit sampling strategies in marine environments, enabling improved characterisation of the spatial genetic structure of dominant reef organisms. Applying this workflow to the weedy coral *Madracis auretenra* on Curaçao, we revealed moderate levels of genotypic diversity despite the species’ predominantly asexual reproductive strategy. Although monospecific stands were typically dominated by a single genotype, substantial genotypic diversity was retained within stands, potentially enhancing ecological resilience under changing environmental conditions. In addition, the detection of a cryptic taxon with evidence of a reproductive barrier suggests additional genetic diversity and potential adaptability in this dominant species. More broadly, our results highlight the potential of photogrammetry-guided sampling to investigate the genetic structure of spatially dominant species in reefscapes where traditional geolocation approaches are limited, providing new opportunities to better understand the adaptive potential and resilience of emerging reef communities.

## Acknowledgements

We would like to acknowledge the “Hope for Reefs Initiative” at the California Academy of Sciences and Inkfish LLC for funding this work. We would like to thank Jade van Dam and Jennifer A. Hoey for their contributions to the data collection during the field surveys. MY was supported by the Aharon and Ephraim Katzir Study Grant from the Israel Academy of Sciences and Humanities, the Murray foundation for student research (2019), and the Alexander von Humboldt Foundation, Germany (2026).

## Data Accessibility and Benefit-sharing

### Data Accessibility

All raw sequences have been uploaded to SRA under the bioproject ID PRJNA1475881. The original code used for data analysis can be found on the GitHub repository associated with this study https://github.com/DennisvHulten/M.auretenra_monostand_diversity.

## Author Contributions

D.v.H. contributed to the conceptualisation of the study and led the data collection, formal analysis, and writing of the paper. M.Y. funded and led the 2021 data collection and photogrammetry processing, and contributed to the conceptualization and editing. M.A.S. contributed to the editing and supervision. L.L. contributed to the analysis, editing, and supervision, and provided part of the funding. P.B. led the conceptualisation, contributed to the analysis and writing, supervised, and provided the primary funding for this study.

