## Supplementary data for "Spatial genotypic patterns within monospecific stands of a dominant coral revealed through photogrammetry-guided sampling"

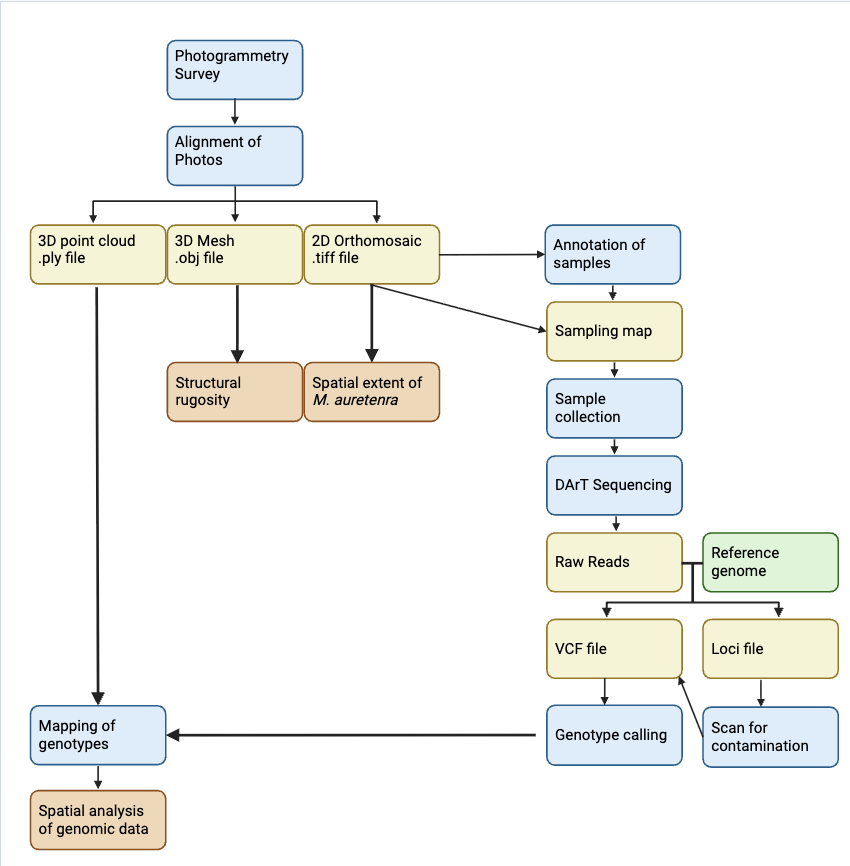

**Figure S1:** Flowchart of the methodology. Processing steps are highlighted in blue, products in yellow, results in orange, and additional resources in green.

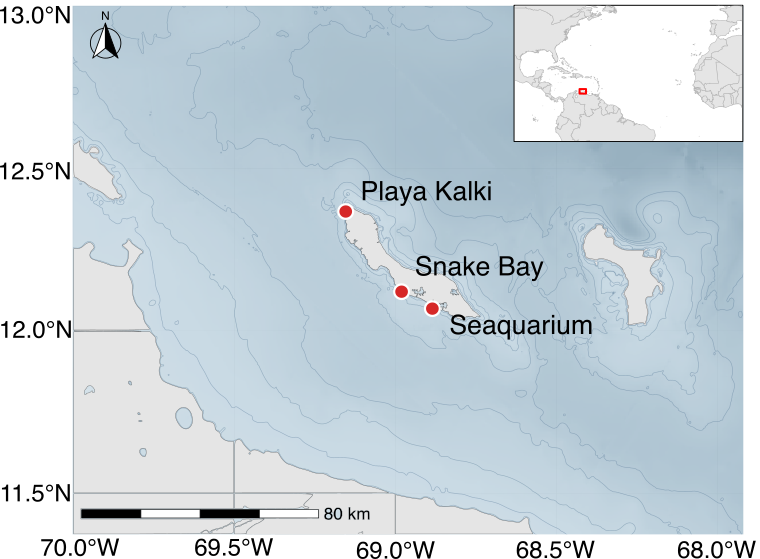

**Figure S2:** Overview of the three sampling sites: Playa Kalki (12.375406, -69.159081), Snake Bay (12.138986, -68.997639), and Seaquarium (12.084367, -68.898439), located on the leeward coast of Curaçao in the Southern Caribbean.

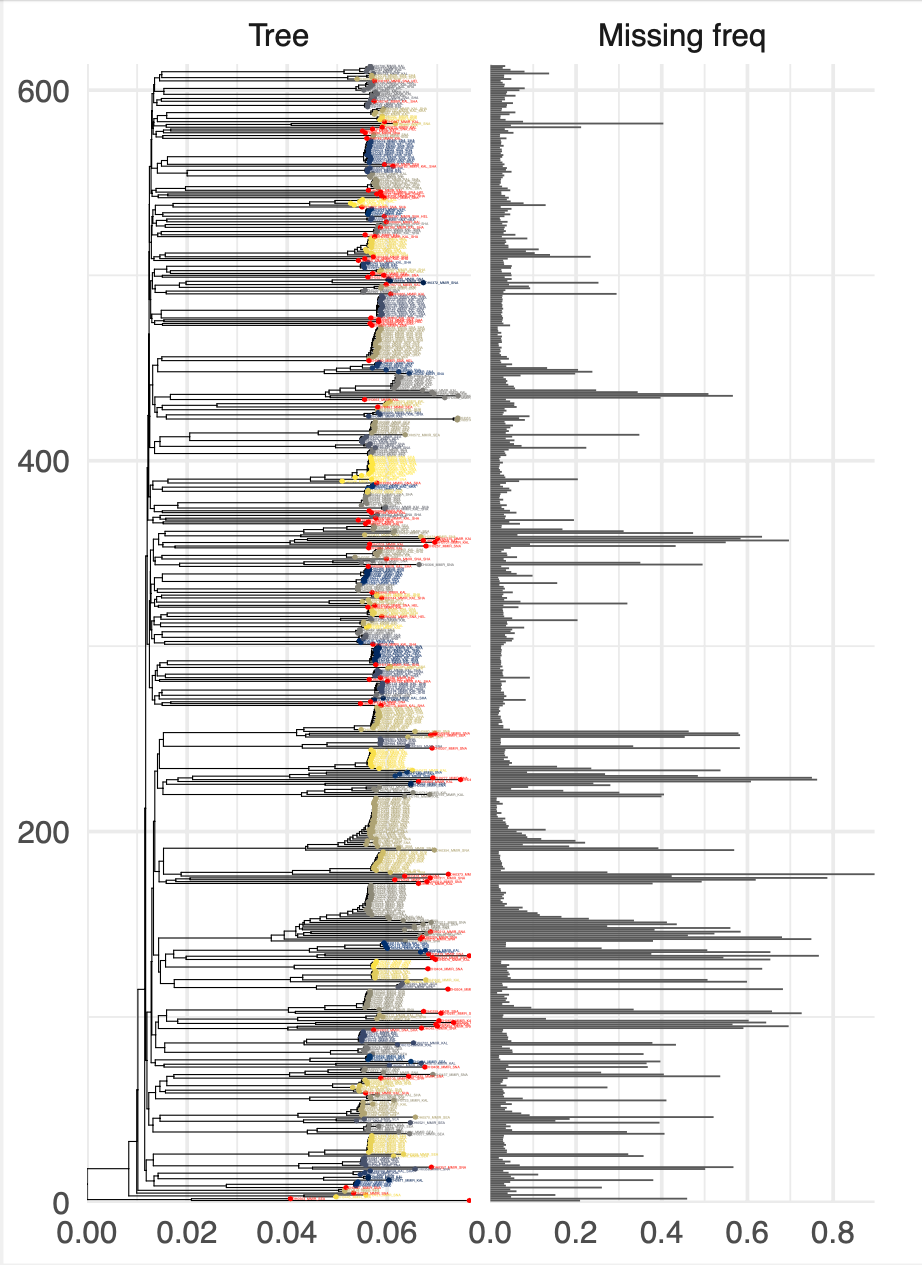

**Figure S3:** Clone detection results based on genetic similarity following a threshold of 98% similarity compared to the percentage of missing data per sample. Clonal groups are highlighted in the same colour, and unique genotypes are highlighted in red.

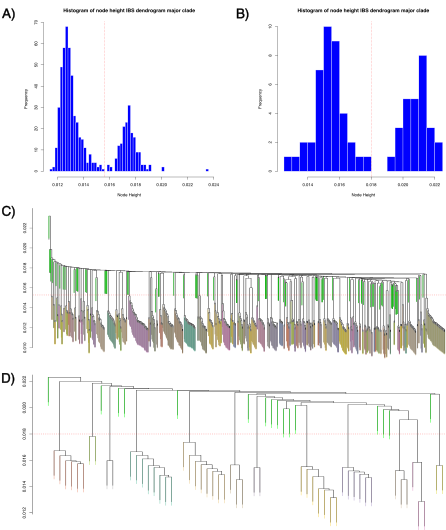

**Figure S4:** Results of the ANGSD IBS distance matrix analysis used to identify clonal groups separated by lineage. A) Histogram of the frequency of node distance between samples of the major lineage. The red dotted line represents the first break between frequencies and is used as the threshold for clonal group detection. B) Histograms of the frequency of node distance between samples of the minor lineage. The red dotted line marks the first break in frequencies and serves as the threshold for clonal group detection. C) IBS distance dendogram of the major lineage with the threshold for clonal groups indicated by the red dotted line. D) IBS distance dendogram of the minor lineage with the threshold for clonal groups indicated by the red dotted line. Distinct colours represent clonal groups, and bright green indicates unique genotypes.

| **Plot** | **Reef zone** | **Lineage** | **Ng** | **N** | **Ng/N** | **Rugosity (2D/3D area)** | **Fractal dimension** |
| --- | --- | --- | --- | --- | --- | --- | --- |
| Snake Bay | Terrace | Major | 10 | 36 | 0.278 | 1.282 | 2.01 |
| Playa Kalki | Terrace | Major | 15 | 32 | 0.469 | 1.401 | 2.02 |
| Playa Kalki | Terrace | Minor | 1 | 1 | 1.000 | 1.401 | 2.02 |
| Snake Bay | Crest | Major | 44 | 149 | 0.295 | 1.479 | 2.09 |
| Playa Kalki | Crest | Major | 61 | 143 | 0.427 | 1.784 | 2.09 |
| Seaquarium | Crest | Major | 2 | 3 | 0.667 | 1.718 | 2.04 |
| Snake Bay | Crest | Minor | 2 | 12 | 0.167 | 1.479 | 2.09 |
| Playa Kalki | Crest | Minor | 1 | 1 | 1.000 | 1.784 | 2.09 |
| Seaquarium | Crest | Minor | 2 | 5 | 0.400 | 1.718 | 2.04 |
| Snake Bay | Slope | Major | 21 | 82 | 0.256 | 1.703 | 2.14 |
| Playa Kalki | Slope | Major | 10 | 30 | 0.333 | 1.785 | 2.14 |
| Seaquarium | Slope | Major | 21 | 79 | 0.266 | 1.791 | 2.06 |
| Snake Bay | Slope | Minor | 12 | 23 | 0.522 | 1.703 | 2.14 |
| Playa Kalki | Slope | Minor | 2 | 3 | 0.667 | 1.785 | 2.14 |
| Seaquarium | Slope | Minor | 5 | 14 | 0.357 | 1.791 | 2.06 |

**Table S1.** Overview of the number of unique multilocus genotypes (Ng), number of sampled colonies (N), genet-to-ramet ratio (Ng/N), and habitat structural complexity metrics (2D/3D rugosity and Fractal dimension) for each site, reef zone, and lineage.

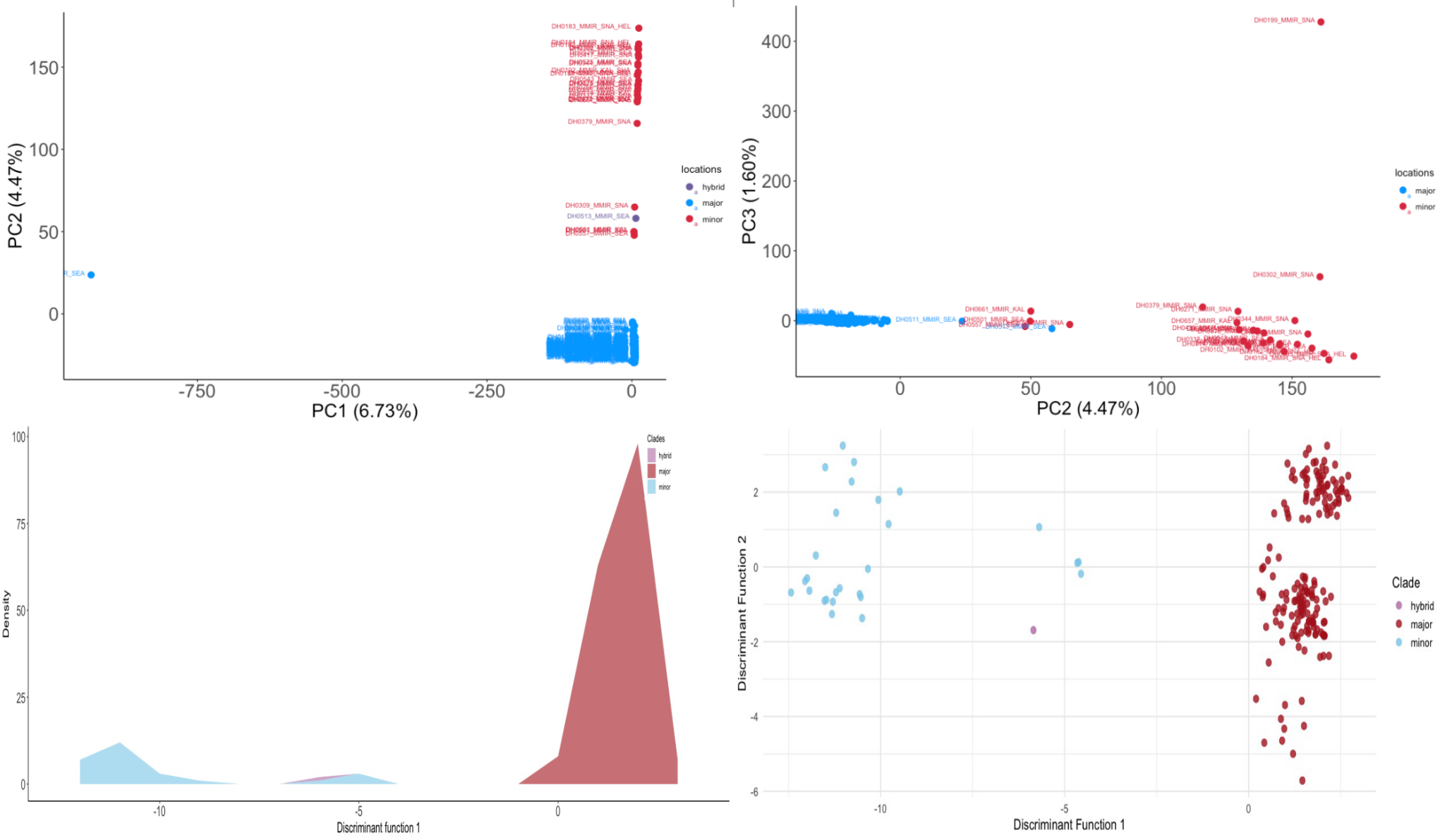

**Figure S5:** Results from the PCA analyses (top) and the denovo DAPC analyses (Bottom).
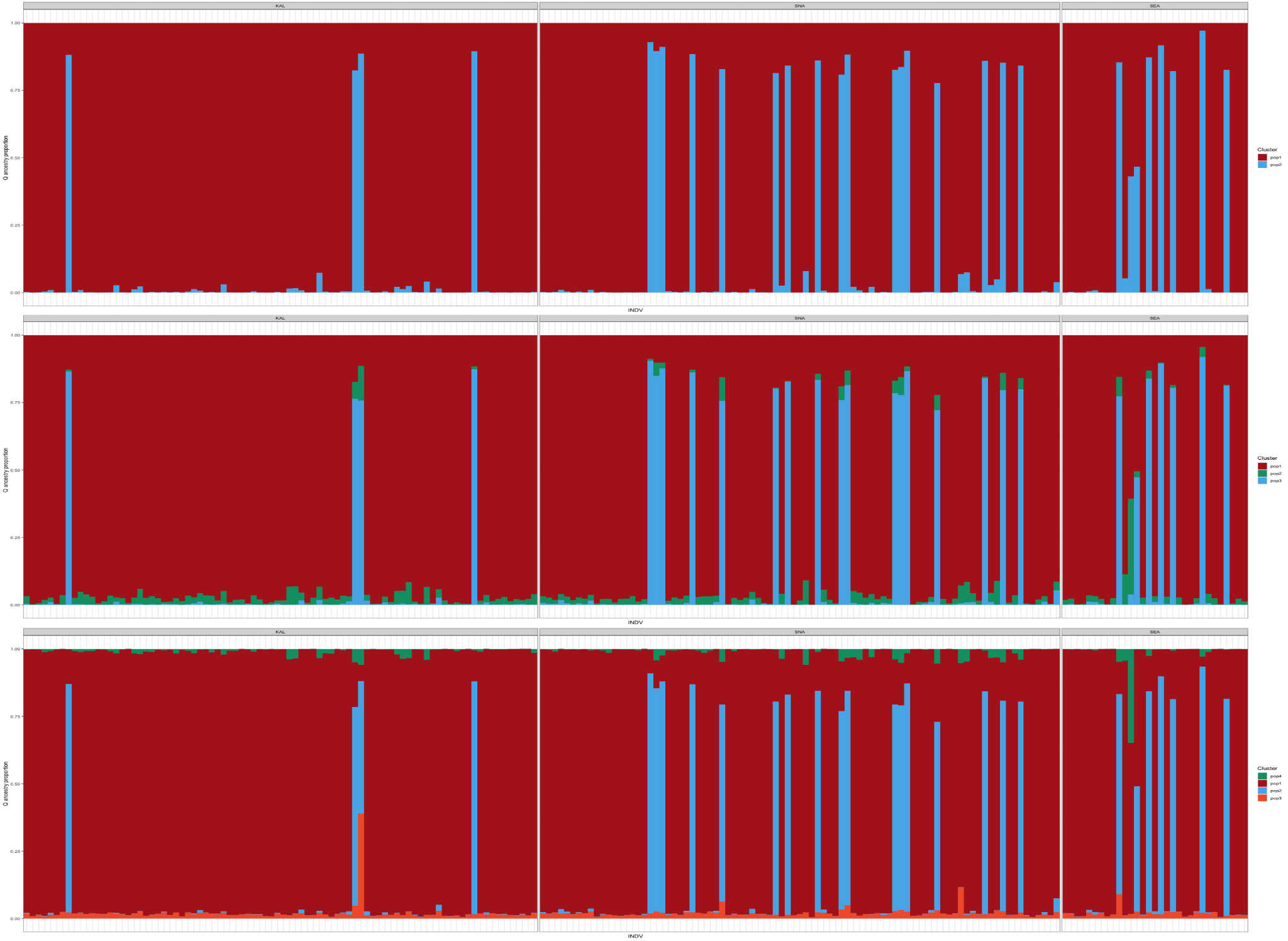

**Figure S6:** Results of the Structure analysis for K = 2, 3, 4. Samples are grouped by site for Playa Kalki (KAL), Snake Bay (SNA), and Seaquarium (SEA).

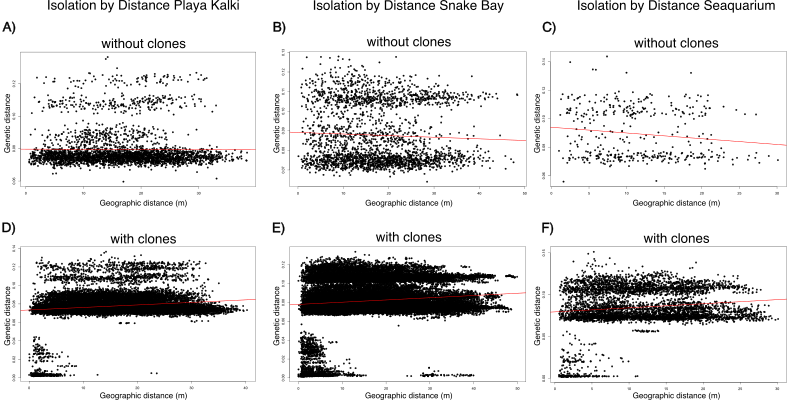

**Figure S7:** Isolation by Distance analysis of unique genotypes across the three sites only including unique genotypes (A,B,C), and including clonal samples (D,E,F). The red line represents a linear regression between geographical distance (m) and genetic distance (Hamming distance).

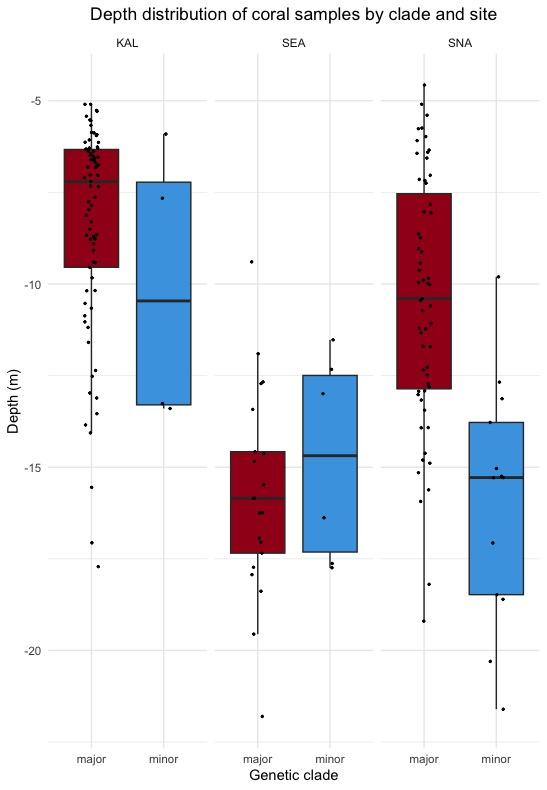

**Figure S8:** Depth distributions of samples from the major and minor lineage, divided by site Playa Kalki (KAL), Seaquarium (SEA), and Snake Bay (SNA).

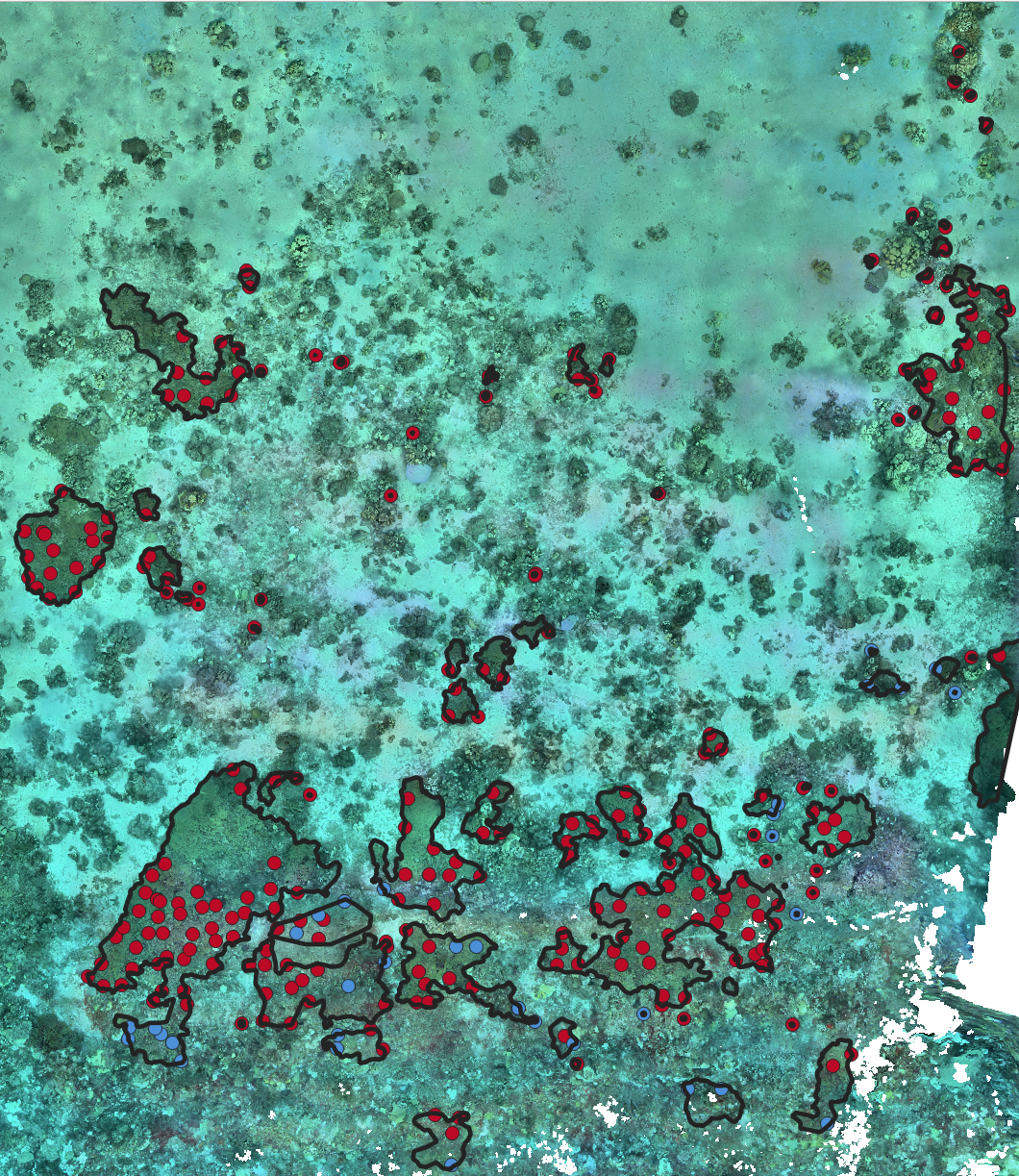

**Figure S9:** Distribution of samples belonging to the major and minor lineages at Snake Bay. Minor lineage members are depicted in blue, whereas major lineage members are depicted in red. Black outlines indicate the extent of colonies, patches and monostands.

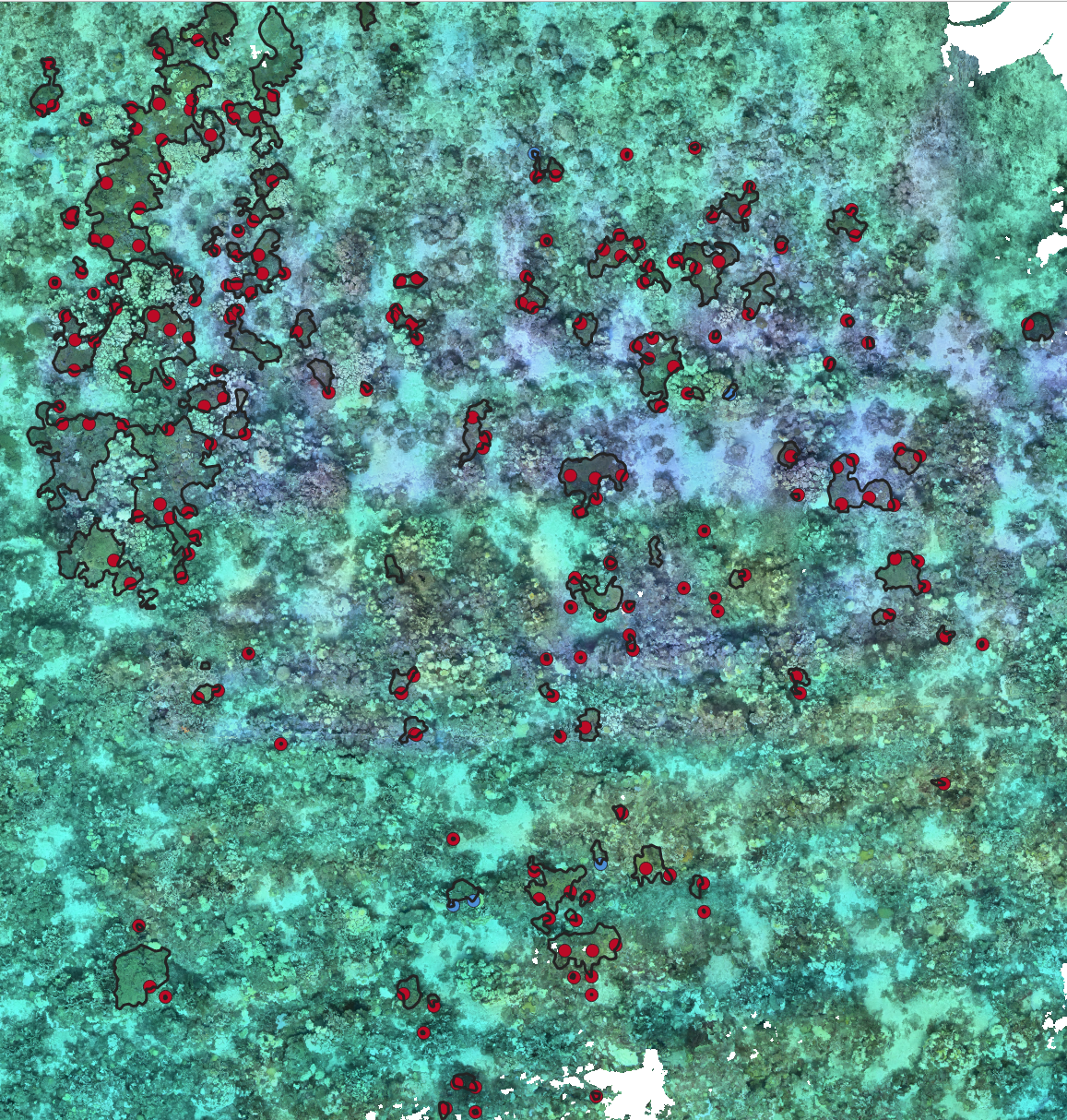

**Figure S10:**  Distribution of samples belonging to the major and minor lineages at Playa Kalki. Minor lineage members are depicted in blue, whereas major lineage members are depicted in red. Black outlines indicate the extent of colonies, patches and monostands.

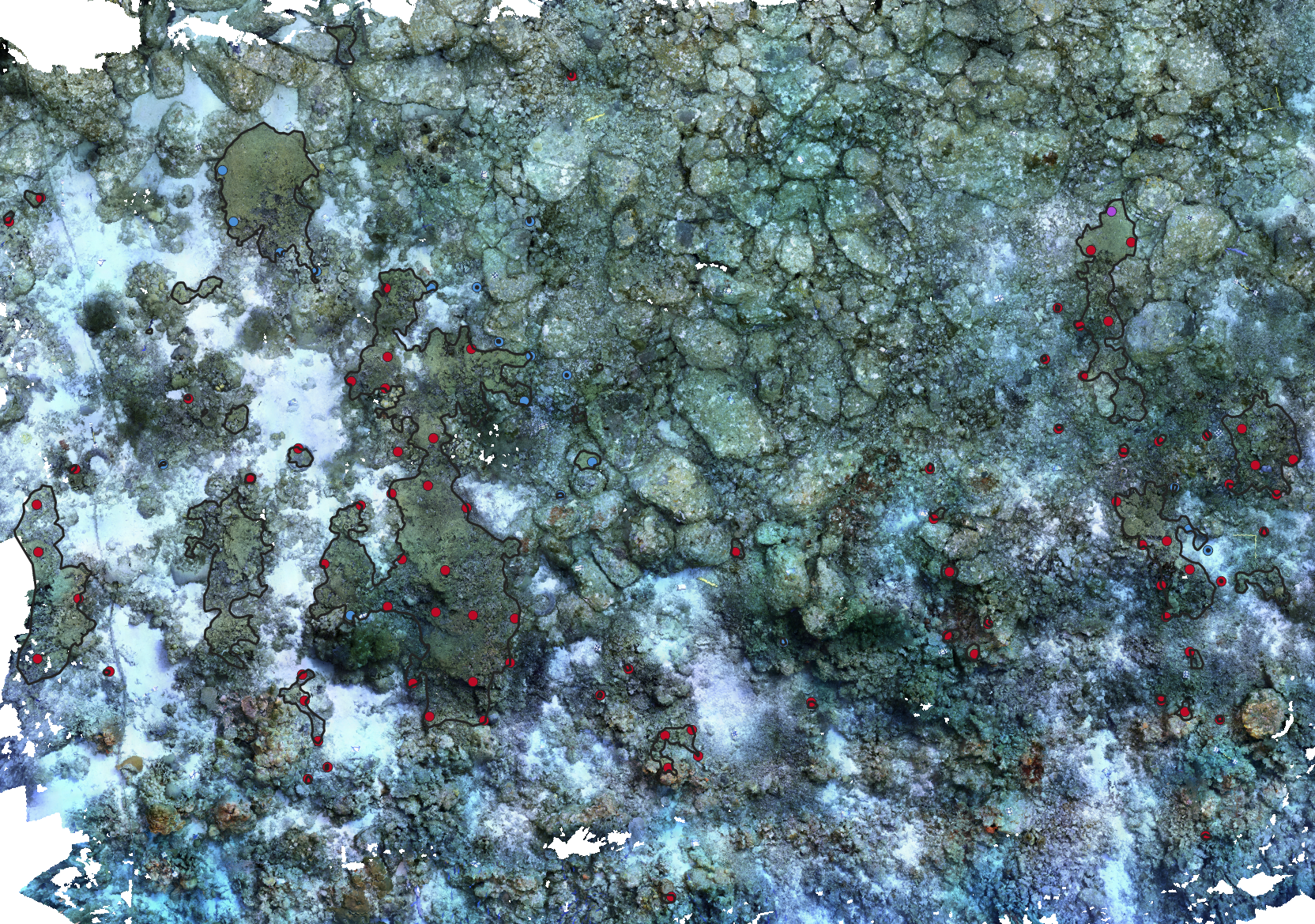
**Figure S11:**  Distribution of samples belonging to the major and minor lineages at Seaquarium. Minor lineage members are depicted in blue, whereas major lineage members are depicted in red. Black outlines indicate the extent of colonies, patches and monostands.

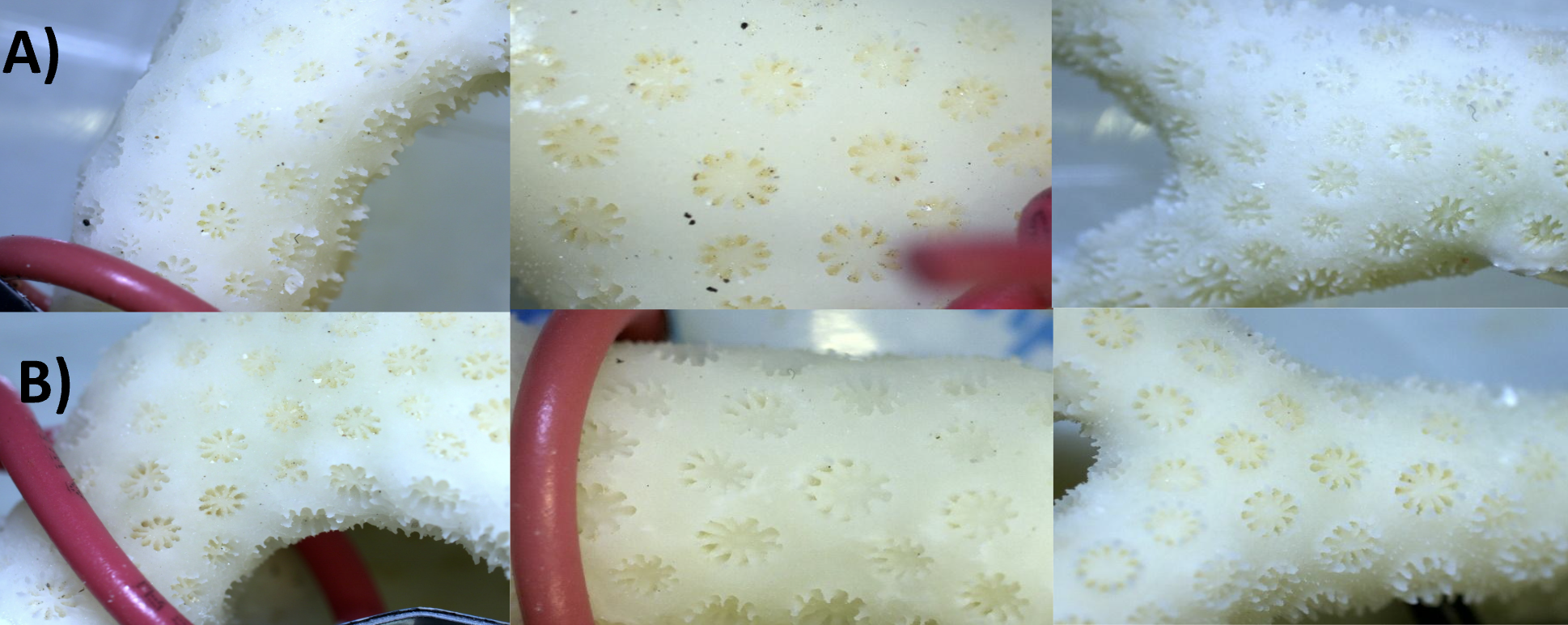

**Figure S12:** Examples of the macrophotographs of skeletal fragments collected for samples of both the major (A) and minor (B) lineage. The number of septa, corallite spacing and corallite density varied both within and across genetic lineages and no evidence for morphological differentiation was observed.

**Table S2:** Combined total cover of *M. auretenra* colonies and aggregations across the three full survey plots and standardised plots.

| Plot | Monospecific stands (m^2^) | Patches (m^2^) | Individual colonies (m^2^) | total cover (m^2^) |
| --- | --- | --- | --- | --- |
| Playa Kalki (full plot) | 36.7 | 29.0 | 19.1 | 84.8 |
| Playa Kalki (standardised plot) | 0.0 | 10.9 | 7.5 | 18.4 |
| Playa Kalki (terrace) | 14.8 | 2.5 | 1.8 | 19.1 |
| Playa Kalki (crest) | 21.9 | 20.1 | 13.6 | 55.6 |
| Playa Kalki (slope) | 0 | 6.4 | 3.7 | 10.0 |
| Snake Bay (full plot) | 145.4 | 17.8 | 9.4 | 172.2 |
| Snake Bay (standardised plot) | 105.0 | 10.6 | 3.3 | 117.9 |
| Snake Bay (terrace) | 3.0 | 0.0 | 1.2 | 4.1 |
| Snake Bay (crest) | 84.2 | 10.0 | 7.0 | 101.2 |
| Snake Bay (slope) | 58.3 | 7.8 | 1.2 | 67.3 |
| Seaquarium (full plot) | 24.9 | 13.4 | 3.7 | 42.0 |
| Seaquarium (standardised plot) | 21.6 | 13.4 | 3.3 | 38.3 |
| Seaquarium (crest) | 4.1 | 0.0 | 0.5 | 4.6 |
| Seaquarium (slope) | 20.8 | 13.4 | 3.2 | 37.3 |

**Table S3:** Structural complexity measurements of all sites based on three different methods: arithmetical mean roughness (Ra), root mean square roughness (Rq), Structural rugosity calculated as 2D vs 3D area (Structural rugosity), and the fractal dimension calculated through the box-counting method (Fractal dimension).

| *Site* | *Environment* | *Ra* | *Rq* | *Structural rugosity* | *Fractal dimension* |
| --- | --- | --- | --- | --- | --- |
| *Playa Kalki* | *Slope* | *0.369* | *0.462* | *1.785* | *2.14* |
| *Playa Kalki* | *Crest* | *0.546* | *0.722* | *1.784* | *2.09* |
| *Playa Kalki* | *Terrace* | *0.207* | *0.259* | *1.401* | *2.02* |
| *Snake Bay* | *Slope* | *0.408* | *0.544* | *1.703* | *2.14* |
| *Snake Bay* | *Crest* | *0.402* | *0.543* | *1.479* | *2.09* |
| *Snake Bay* | *Terrace* | *0.140* | *0.198* | *1.282* | *2.01* |
| *Seaquarium* | *Slope* | *0.337* | *0.431* | *1.791* | *2.06* |
| *Seaquarium* | *Crest* | *0.525* | *0.647* | *1.718* | *2.04* |
| *Seaquarium* | *Terrace* | *NA* | *NA* | *NA* | *NA* |

**Table S5:** Percentage shared genotypes between sampling groups.

| Source 1 | Source 2 | Total from | Shared | Shared % |
| --- | --- | --- | --- | --- |
| patch | monostand | 77 | 22 | 28.57143 |
| isolated | monostand | 78 | 29 | 37.17949 |
| monostand | patch | 98 | 22 | 22.44898 |
| isolated | patch | 78 | 36 | 46.15385 |
| monostand | isolated | 98 | 29 | 29.59184 |
| patch | isolated | 77 | 36 | 46.75325 |

**Table S6:** Simulation results for sampling 20 random samples from each sampling group.

| Sampling group | Probability of only unique genotypes | Average number of unique genotypes |
| --- | --- | --- |
| Monospecific stands | 0.00351 | 15.5 |
| Monospecific stands + Isolated colonies | 0.0554 | 17.4 |
| Monospecific stands + Patches | 0.0616 | 17.5 |
| Patches | 0.0563 | 17.5 |
| Monospecific stands + Isolated colonies + Patches | 0.116 | 18.0 |
| Isolated colonies + Patches | 0.130 | 18.1 |
| Isolated colonies | 0.126 | 18.2 |

**Table S7:** Simulation results for sampling 20 random samples from each sampling group.

| Sampling group | Site | Probability of only unique genotypes | Average number of unique genotypes |
| --- | --- | --- | --- |
| isolated | Playa Kalki | 0.02855 | 17.16174 |
| monostand+isolated+patch | Playa Kalki | 0.02678 | 16.87841 |
| isolated+patch | Playa Kalki | 0.01915 | 16.61503 |
| monostand+isolated | Playa Kalki | 0.01257 | 16.44508 |
| monostand+patch | Playa Kalki | 0.00601 | 16.01373 |
| patch | Playa Kalki | 0.00216 | 15.46171 |
| monostand | Playa Kalki | 0.00000 | 10.25820 |
| monostand+isolated+patch | Seaquarium | 0.00003 | 13.83935 |
| monostand+isolated | Seaquarium | 0.00002 | 13.55457 |
| isolated | Seaquarium | 0.00000 | 13.33260 |
| monostand+patch | Seaquarium | 0.00000 | 13.10183 |
| isolated+patch | Seaquarium | 0.00000 | 13.02622 |
| patch | Seaquarium | 0.00000 | 10.26237 |
| monostand | Seaquarium | 0.00000 | 10.00570 |
| monostand+isolated+patch | Snake Bay | 0.00215 | 15.22285 |
| isolated+patch | Snake Bay | 0.00104 | 15.06710 |
| monostand+patch | Snake Bay | 0.00089 | 14.72191 |
| monostand+isolated | Snake Bay | 0.00094 | 14.62664 |
| isolated | Snake Bay | 0.00009 | 14.44878 |
| patch | Snake Bay | 0.00003 | 14.43894 |
| monostand | Snake Bay | 0.00004 | 13.51536 |
